# *Unraveling Amazon tree community assembly using Maximum Information Entropy*: a quantitative analysis of tropical forest ecology

**DOI:** 10.1101/2021.03.31.437717

**Authors:** Edwin Pos, Luiz de Souza Coelho, Diogenes de Andrade Lima Filho, Rafael P. Salomão, Iêda Leão Amaral, Francisca Dionízia de Almeida Matos, Carolina V. Castilho, Oliver L. Phillips, Juan Ernesto Guevara, Marcelo de Jesus Veiga Carim, Dairon Cárdenas López, William E. Magnusson, Florian Wittmann, Mariana Victória Irume, Maria Pires Martins, Daniel Sabatier, José Renan da Silva Guimarães, Jean-François Molino, Olaf S. Bánki, Maria Teresa Fernandez Piedade, Nigel C.A. Pitman, Abel Monteagudo Mendoza, José Ferreira Ramos, Joseph E. Hawes, Everton José Almeida, Luciane Ferreira Barbosa, Larissa Cavalheiro, Márcia Cléia Vilela dos Santos, Bruno Garcia Luize, Evlyn Márcia Moraes de Leão Novo, Percy Núñez Vargas, Thiago Sanna Freire Silva, Eduardo Martins Venticinque, Angelo Gilberto Manzatto, Neidiane Farias Costa Reis, John Terborgh, Katia Regina Casula, Euridice N. Honorio Coronado, Juan Carlos Montero, Beatriz S. Marimon, Ben Hur Marimon-Junior, Ted R. Feldpausch, Alvaro Duque, Chris Baraloto, Nicolás Castaño Arboleda, Julien Engel, Pascal Petronelli, Charles Eugene Zartman, Timothy J. Killeen, Rodolfo Vasquez, Bonifacio Mostacedo, Rafael L. Assis, Jochen Schöngart, Hernán Castellanos, Marcelo Brilhante de Medeiros, Marcelo Fragomeni Simon, Ana Andrade, José Luís Camargo, Layon O. Demarchi, William F. Laurance, Susan G.W. Laurance, Emanuelle de Sousa Farias, Maria Aparecida Lopes, José Leonardo Lima Magalhães, Henrique Eduardo Mendonça Nascimento, Helder Lima de Queiroz, Gerardo A. Aymard C., Roel Brienen, Juan David Cardenas Revilla, Flávia R.C. Costa, Adriano Quaresma, Ima Célia Guimarães Vieira, Bruno Barçante Ladvocat Cintra, Pablo R. Stevenson, Yuri Oliveira Feitosa, Joost F. Duivenvoorden, Hugo F. Mogollón, Leandro Valle Ferreira, James A. Comiskey, Freddie Draper, José Julio de Toledo, Gabriel Damasco, Nállarett Dávila, Roosevelt García-Villacorta, Aline Lopes, Alberto Vicentini, Janaína Costa Noronha, Flávia Rodrigues Barbosa, Rainiellen de Sá Carpanedo, Thaise Emilio, Carolina Levis, Domingos de Jesus Rodrigues, Juliana Schietti, Priscila Souza, Alfonso Alonso, Francisco Dallmeier, Vitor H.F. Gomes, Jon Lloyd, David Neill, Daniel Praia Portela de Aguiar, Alejandro Araujo-Murakami, Luzmila Arroyo, Fernanda Antunes Carvalho, Fernanda Coelho de Souza, Dário Dantas do Amaral, Kenneth J. Feeley, Rogerio Gribel, Marcelo Petratti Pansonato, Jos Barlow, Erika Berenguer, Joice Ferreira, Paul V.A. Fine, Marcelino Carneiro Guedes, Eliana M. Jimenez, Juan Carlos Licona, Maria Cristina Peñuela Mora, Carlos A. Peres, Boris Eduardo Villa Zegarra, Carlos Cerón, Terry W. Henkel, Paul Maas, Marcos Silveira, Juliana Stropp, Raquel Thomas-Caesar, Tim R. Baker, Doug Daly, Kyle G. Dexter, John Ethan Householder, Isau Huamantupa-Chuquimaco, Toby Pennington, Marcos Ríos Paredes, Alfredo Fuentes, José Luis Marcelo Pena, Miles R. Silman, J. Sebastián Tello, Jerome Chave, Fernando Cornejo Valverde, Anthony Di Fiore, Renato Richard Hilário, Juan Fernando Phillips, Gonzalo Rivas-Torres, Tinde R. van Andel, Patricio von Hildebrand, Edelcilio Marques Barbosa, Luiz Carlos de Matos Bonates, Hilda Paulette Dávila Doza, Émile Fonty, Ricardo Zárate Gómez, Therany Gonzales, George Pepe Gallardo Gonzales, Jean-Louis Guillaumet, Bruce Hoffman, André Braga Junqueira, Yadvinder Malhi, Ires Paula de Andrade Miranda, Linder Felipe Mozombite Pinto, Adriana Prieto, Agustín Rudas, Ademir R. Ruschel, Natalino Silva, César I.A. Vela, Vincent Antoine Vos, Egleé L. Zent, Stanford Zent, Bianca Weiss Albuquerque, Angela Cano, Diego F. Correa, Janaina Barbosa Pedrosa Costa, Bernardo Monteiro Flores, Milena Holmgren, Marcelo Trindade Nascimento, Alexandre A. Oliveira, Hirma Ramirez-Angulo, Maira Rocha, Veridiana Vizoni Scudeller, Rodrigo Sierra, Milton Tirado, Maria Natalia Umaña, Geertje van der Heijden, Emilio Vilanova Torre, Corine Vriesendorp, Ophelia Wang, Kenneth R. Young, Manuel Augusto Ahuite Reategui, Cláudia Baider, Henrik Balslev, Sasha Cárdenas, Luisa Fernanda Casas, William Farfan-Rios, Cid Ferreira, Reynaldo Linares-Palomino, Casimiro Mendoza, Italo Mesones, Armando Torres-Lezama, Ligia Estela Urrego Giraldo, Daniel Villarroel, Roderick Zagt, Miguel N. Alexiades, Karina Garcia-Cabrera, Lionel Hernandez, William Milliken, Walter Palacios Cuenca, Susamar Pansini, Daniela Pauletto, Freddy Ramirez Arevalo, Adeilza Felipe Sampaio, Elvis H. Valderrama Sandoval, Luis Valenzuela Gamarra, Gerhard Boenisch, Jens Kattge, Nathan Kraft, Aurora Levesley, Karina Melgaço, Georgia Pickavance, Lourens Poorter, Hans ter Steege

**Author notes:** Deceased 01-2018 (Jean-Louis Guillaumet).

## Abstract

In a time of rapid global change, the question of what determines patterns in species abundance distribution remains a priority for understanding the complex dynamics of ecosystems. The constrained maximization of information entropy provides a framework for the understanding of such complex systems dynamics by a quantitative analysis of important constraints via predictions using least biased probability distributions. We apply it to over two thousand hectares of Amazonian tree inventories across seven forest types and thirteen functional traits, representing major global axes of plant strategies. Results show that constraints formed by regional relative abundances of genera explain eight times more of local relative abundances than constraints based on directional selection for specific functional traits, although the latter does show clear signals of environmental dependency. These results provide a quantitative insight by inference from large-scale data using cross-disciplinary methods, furthering our understanding of ecological dynamics.

## Introduction

Drivers of species distributions and their predictions have been a long-standing search in ecology, with approaches varying from deterministic to neutral (i.e. stochastic) and almost everything in between (e.g. near-neutral, continuum or emergent-neutral [1,2]). Most models are based on prior assumptions of processes that drive community dynamics. The Maximum Entropy Formalism (hereafter called MEF) makes no such, potentially unjustified, a-priori assumptions in generating predictions of species abundance distributions, as such it is a useful construct to infer processes driving community dynamics given the constraints imposed by prior knowledge (e.g. functional traits or summed regional abundances) [3]. Quantifying the relative importance of these distinct constraints can thus provide additional answers to understand the complexity of community dynamics (see Supporting Materials SM: boxes S1-S3). This is especially so because, although many different tests are available that link variation in taxon abundances to 1) trait variation, 2) taxon turnover between habitats or environments and 3) the distance decay of similarities between samples, none quantify the importance of these relative to each other. The MEF as applied here, however, is capable of and designed to do exactly this by decomposing variation to separate information explained by each of these aspects in a four-step model (Fig. 1 and Box S2). Its application to an unprecedented large tree inventory database on genus level taxonomy consisting of > 2,000 1-ha plots distributed over Amazonia [4] and a genus trait database of 13 key functional traits representing global axes of plant strategies [5] allows us to advance the study of Amazonian tree community dynamics from a new cross-disciplinary perspective.

**Fig. 1.**
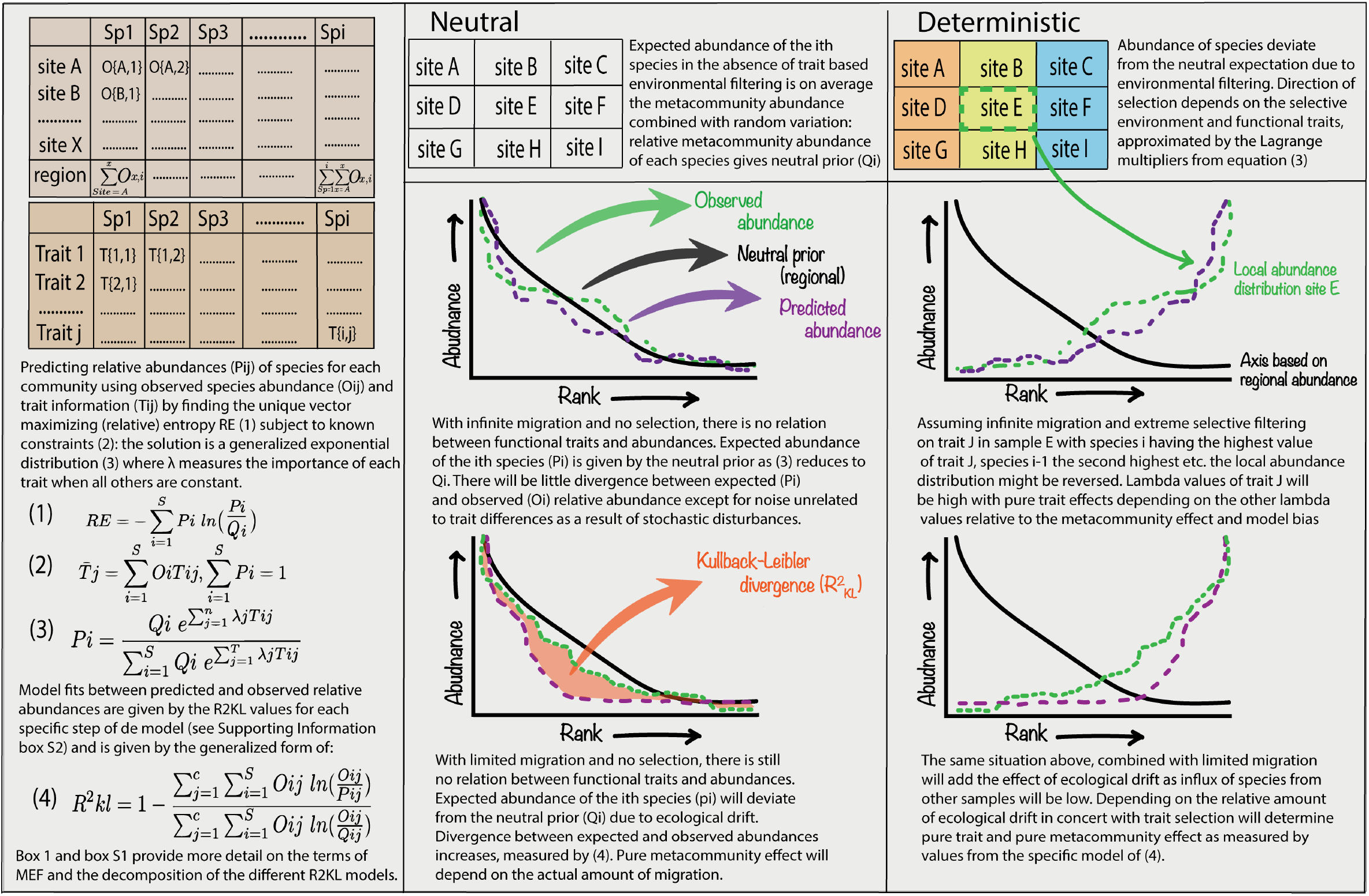
Schematic depiction of the MEF procedure. Left panel shows a genus abundances per site and a functional trait matrix per genus, bottom half outlines calculations. Middle and right panel show different scenarios of neutral and deterministic dynamics under infinite or limited migration. Figure was custom made using Adobe Illustrator (Adobe Inc., 2019. Adobe Illustrator).

## Results

Principles from information theory [6–8] can be used in an ecological setting to predict the most likely abundance state for each taxon while simultaneously maximizing entropy based on constraints. Maximization of entropy allows quantifying the information yield for each constraint and therefor identifies which constraints reduce entropy the most. Here we specifically use Shipley’s mathematical framework (CATS) for the MEF calculations, similar to earlier studies [9–11].

### Predictive power of the four-step model

Using a uniform prior and CWM values (Community Weighted Means) as constraints accounted for 23% on average of total deviance between observed and predicted relative abundances (measured by R^2^_KL_ values, see Box S2 equation 5). Filtered by forest type this was 34% for podzol forests, *várzea* 25%, *igapó* 23%, swamp forests 34%, 21% and 24% for Guyana Shield and Pebas *terra firme* respectively and 20% for Brazilian Shield *terra firme* forests (see Table S1 for detailed decomposition).

Using observed metacommunity relative abundances as prior regardless of functional traits accounted for on average 56% for the combined dataset with for all forest types between 51 and 55%, except for the Guyana Shield *terra firme* with 62%. The hybrid model (including both traits as constraints and the metacommunity prior) performed slightly better for the combined dataset (average 60%) with a minimum of 57% for swamp and *várzea* forests and a maximum of 66% for Guyana Shield *terra firme* forests. To compensate for spurious relationships between regional abundances and local trait constraints, regardless of selection, explanatory power was regarded relative to model bias yielding the pure trait and metacommunity effects (Box S3, Fig. 2 and Table S1). This lowered the proportion of information accounted for and yielded average pure metacommunity effects of 40% for the overall dataset ranging between 26 and 45% for each forest type separately with pure trait effects explaining only 5% of information for the combined dataset on average with for each forest type between 3 and 9%. Although the latter was lowered substantially, the explanatory power did appear to be strongly dependent on forest type. The online supplementary material provides additional results relating to the predictive power of each model as well as the spatial gradient of the pure trait and metacommunity effects (Figs. S2-3).

**Fig. 2.**
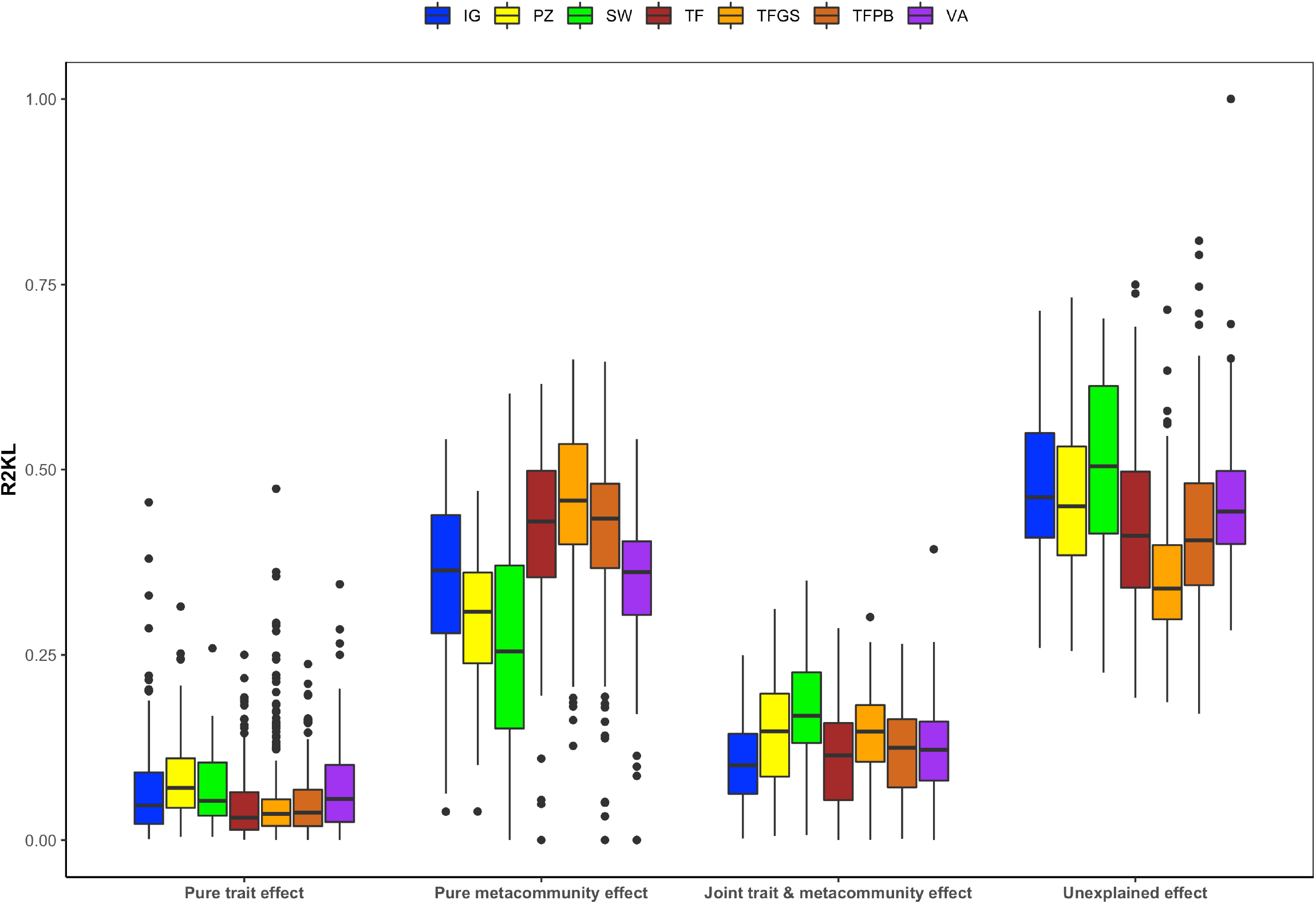
Visual representation of pure trait, pure metacommunity, hybrid model and the remaining unexplained information for each separate forest type. Abbreviations indicate different types: igapó (IG), podzol (PZ), swamp (SW), Brazilian shield terra firme (TFBS), Guiana Shield terra firme (TFGS), Pebas terra firme (TFPB) and várzea (VA). Boxplots show median value of pure effects over all samples, with lower and upper hinges corresponding to 25th and 75th percentiles. Whiskers extends from hinge to largest or smallest value no further than 1.5 * IQR from hinge. Points beyond this range are plotted individually and only positive values were plotted.

### Direction and strength of selection of trait-based constraints

Each trait showed significant differences in lambda when compared between forest types (Fig. S1, see methods for a definition of lambda). Scatterplots of CWM trait values versus lambda show that, in general, higher lambda values correspond with higher CWM trait values (Figure S7). A number of functional traits associated with low nutrient conditions (e.g. ectomycorrhiza) and life history strategies suited for protection against herbivores (e.g. latex, resin and high leaf C content) were clearly positively associated with abundance in nutrient poor environments (podzols), indicated by the positive lambda values. In contrast, having fleshy fruits and high leaf N and P content were clearly negatively associated with abundance on these soils.

Nodulation was also negatively associated with abundance on poor soils. The ability to accumulate aluminium was positively associated with abundance on those soils commonly associated with higher aluminium content such as *igapó* (strong positive effects). In contrast, it was strongly negatively associated with abundance on other soils, with negative lambda values for podzol and *várzea* forests. Traits such as SLA or winged fruits also showed patterns dependent on forest type, although less pronounced.

### Effect of regional metacommunity prior

There was a remarkable similar mean 22% decrease of the information explained purely by the metacommunity prior for each forest type (Fig. S3). For the separate forest types, although the initial pure metacommunity effect varied, the decline appeared remarkably similar with a mean 25% decrease in pure metacommunity effect for podzol, 23% for *várzea*, 26% for *igapó* and 27% for swamp forests with *terra firme* forests having a smaller decline of approximately 21%, averaged over the three subregions. It should be noted there is an obvious risk that when sampling size is increased, this also includes more environmental heterogeneity as samples are coming from a variety of localities potentially leading to changing composition. If this were the case, however, the regional prior (q_i_ from Fig. 1 and Box S2) would also change, as taxa might be abundant in some places but rare or absent in others. As the metacommunity effect is the explained information that remains relative to any trait effects (i.e. information unique to the neutral prior) and the pure trait effects are the explained information remaining after correcting for pure metacommunity effects (Box S3) this effect should then be accompanied by an increase in pure trait effect for each sample. This was not observed, not even within the different forest types.

Instead, the trait effect gradually went up and then remained constant (Fig. S4).

## Discussion

The MEF emerges from a well-founded theoretical and empirical body of ecology and evolutionary biology, regarding natural selection, migration and population dynamics. From an ecological point of view, it can be used to quantify the relative association between directional or stabilizing selection for functional traits versus the importance of relative regional abundance regardless of these traits by imposing these as constraints. Our results show that pure trait effects, on average, explained only 5% of the information when all forest types were taken together whereas the pure metacommunity effect, however, explained eight times more taken all forest types together (40%). Greater trait dissimilarity was positively associated with higher pure trait effects, indicating trait-based selection, although the assumed influence of dispersal regardless of these traits appeared to confer more information explaining tree genus composition of the Amazon rainforest. The strength and direction of selection indicated clear directional selective pressure for life history strategies of either growth or protection, depending on forest type (see supplementary online material S-A for a more detailed exploration of ecological interpretation). Including community weighted variance as reflective of potential stabilizing selection did not provide additional information. Although this could be interpreted as indicative of weak or absent stabilizing selection, it is more likely to be an artefact of many genera not being shared among localities due to the sheer geographical scale resulting in a strong mismatch between observed, predicted and uniform relative abundances resulting in a model bias higher than information yielded by including these constraints (see also box S2).

Despite showing clear patterns in environmental selection and dispersal effects, there was a large proportion of information left unexplained (44% on average). Potentially, local demographic stochasticity could weaken any link between functional traits measured and regional abundances of genera. This would, however, mean that almost half of the information contained in relative abundances are the result of random population dynamics and are not structurally governed. Alternatively, this could be due to functional traits reflective of processes not taken into account in this study, such as traits reflective of interactions between trophic levels (e.g. traits influencing specific plant-pathogen interactions). Another and at least equally likely hypothesis for (local) unexplained information is that when scaling up, the ratio of genus richness to total abundance decreases rapidly initially but stabilizes again as relatively non-overlapping habitats are included in the regional abundance distributions and more genera are included again due to the different habitats. This would result in a change of the regional abundance distribution (i.e. the prior) to which each local community is compared, resulting in higher local unexplained information. Further study into these aspects could provide additional insight, though the data necessary for these scales is lacking for Amazonian trees.

### Metacommunity importance

Although the initial explanatory power of the metacommunity prior differed between forest types, the decay pattern was very similar. As the effects of either traits or the metacommunity are measured in the goodness-of-fit predictions on local relative abundances, this implies that at small spatial scales the surrounding regional abundances provide better estimators than functional traits, while at larger spatial scales this shifts to the traits. The ecological translation would be that on small spatial scales, local communities share similar environmental conditions leaving dispersal and drift acting predominantly in changing community composition, at least for genus level taxonomy. As the potential regional pool is increased, more and more environmental heterogeneity and non-overlapping regions are likely to be introduced. The more gradual decline of *terra firme* forests can then arguably be attributed to these forests having the largest relative surface area of Amazonia (even for the separate subregions), potentially giving these forests an almost continuous metacommunity without gaps, resulting in a more gradual transition from metacommunity to trait relative importance. The fact that metacommunity effects do not change anymore after certain distances would indicate the effect of dispersal potentially occurs over very large distances. It should be noted that as these calculations are done at community and genus level, they do not measure single dispersal events but rather the effect of dispersal on community composition much deeper in time. In other words, this effect suggests more than a dispersal event every now and then. Instead, it argues for prolonged mixing of forests on large geographical and temporal scales, supported by recent findings demonstrating a lack of geographical phylogenetic structure of lineages for Amazonian tree genera [12].

### Conclusion

Using an unprecedented scale of data and applying the Maximum Entropy Formalism from information theory we show that constraints formed by regional relative abundances of genera explain eight times more of local relative abundances then constraints based on directional selection for specific functional traits, although the latter does show clear signals of environmental dependency. There is, however, still much to be explored due to the large unexplained effects and analyses on finer taxonomic (i.e. species level) and environmental (e.g. microhabitat) scales could resolve these issues. The relatively large effects of the regional pool of genera over great distances does suggest an important role for long term dispersal and mixing of Amazonian trees, especially for the Amazonian interior.

## Methods

### Empirical data

The ATDN [4,13,14] consists of over 2000 tree inventory plots distributed over the Amazon basin and the Guiana Shield, collectively referred to as Amazonia (a map of all current plots can be found at https://atdn.myspecies.info/). Only those plots with trees ≥ 10 cm diameter at breast height were used, leaving 2011 plots with a mean of 558 individuals per plot identified to at least genus level. Most plots used are 1 ha in size (1414) with 492 being smaller (minimum size of .1 ha) and 105 larger (maximum size of 80 ha). Genera have been standardized to the W3 Tropicos database [15] using the Taxonomic Name Resolution Service (TNRS, see [16]). After filtering based on above criteria and solving nomenclature issues, 1,121,935 individuals belonging to over 828 genera remained. Plots were distributed over seven abiotically different forest types: Podzol forests (PZ), *Igapó* (IG, black water flood forests), *Várzea* (VA white water flood forests), Swamp (SW) and *Terra firme* forests (TF) with subregions BS (Brazilian Shield), GS (Guyana Shield) and PB (Pebas) (see also [17] for details regarding these forest types). Trait data were extracted from several sources. Wood density was mostly derived from [18]. Traits related to leaf characteristics mostly came from four large datasets [19–24], including additional data from other sources [25–27] as well as unpublished data (J. Lloyd, A.A. de Oliveira, L. Poorter, M. van de Sande & Mazzei, M. van de Sande & L. Poorter). Data on seed mass came from [28–30] as well as different flora’s and tree guides. As this particular trait can vary over several orders of magnitude, this was included on a log-scale [29,31].

Ectomycorrhizal aspects were derived from literature [32], the same applies to nodulation [33,34]. Traits involved in aluminum accumulation were based on [35,36] and references therein. For binary traits (yes/no), a genus was considered having a certain trait only when >50% of the genus was positive for that specific trait.

### Functional traits and trait imputation

Constraints were formed by Community Weighted genus Means (CWM) of functional traits (Table 1), related to key ecological life history aspects. According to principles of natural selection, CWM values will likely be biased towards favourable trait values for that particular environment in the case of directional selection, as taxa with these traits will be more abundant due to environmental selection. Previous studies included community weighted variance (CWV) as well as indicative of potential stabilizing selection [11,37]. In our case, however, including CWV as constraints resulted in a model bias that was consistently higher than information including trait or metacommunity aspects, CWV was therefore not included as constraints in the final analysis. As for many traits it has been shown earlier that the interspecific variability was larger than the intraspecific variability, this allowed the use of data from different sources to at least calculate a mean species trait value. Genus trait values were subsequently computed as genus-level means of species values if known within the genus and considered constant for each genus. Genus level of taxonomy was used as the available trait database had the most information on this taxonomic level (see Table 1). Unknown values for traits were estimated by Multiple Imputation with Chained Equations (MICE, see [15]) by delta adjustment, subtracting a fixed amount (delta), with sensitivity of this adjustment to the imputations of the observed versus imputed data analysed using density plots (Fig. S8) and a linear regression model. This procedure was done using the *mice* package [38], available on the R repository, under predictive mean matching (*pmm* setting, 50 iterations). Results showed imputations were stable and showed near identical patterns with each imputation scenario (see Figs S5-6 and Table S2). After imputation, all trait values were transformed to Community Weighted Means (CWM) of each trait (*J*) for each plot 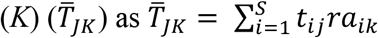 with *ra* the relative abundance of the *i^th^* genus in the *k^th^* plot following earlier uses [37].

**Table 1.**
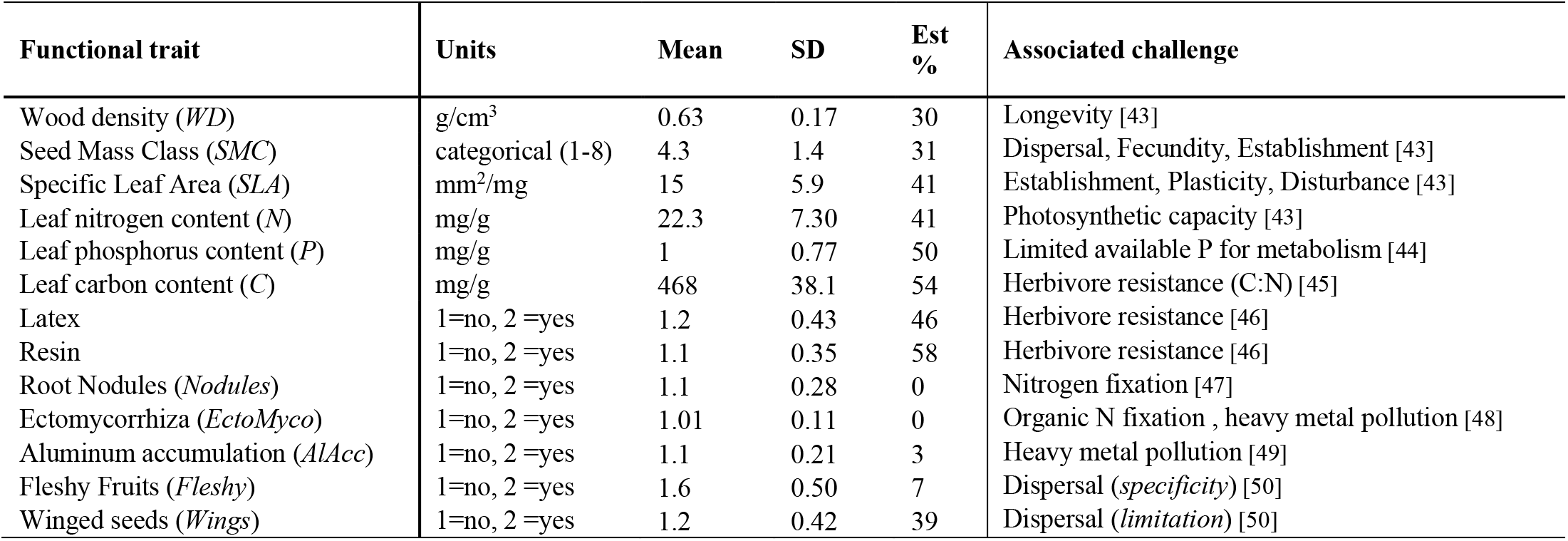
Overview of used functional traits. Mean and standard deviation (SD) are calculated after predictive mean matching (percentage of estimated values is given by Est (%)). Associated challenge indicates different aspects of life history and selective environment related to specific functional traits, sources are given in the footnote. For specific methodology of measurement protocols and calculation for each trait we refer to the original sources of the data (see main text).

### MEF procedure predictions and ecological inference

Figure 1 provides a schematic procedure overview, box S1 provides an overview of important terms and Boxes S2-3 further mathematical details. Initially, a maximally uninformative prior is specified, where q_i_ (Box S1 equation 1) equals 1/S, indicative of each species having equal abundances, and trait constraints are randomly permuted multiple times (n=50) among genera to test whether inclusion of specified constraints significantly changes derived probability distributions (see also [39]). Subsequently, the same prior is used but now observed trait CWM values belonging to specific genera are used as constraints (following earlier applications using simulated communities [11]). Third, observed regional abundances are used as prior with permutated trait constraints and finally both observed regional abundances and observed trait CWM are used as prior and constraints. *Maxent2*, an updated version of the *maxent* function currently in the FD library of R provided the computational platform. Proportions of uncertainty explained by each model are given by the Kullback-Leibler divergence R^2^_KL_, a generalization of the classic R^2^ goodness of fit.

In contrast with standard linear regression models having squared goodness-of-fits measurements, the R^2^_KL_ is much more related to the concept of relative entropy, quantifying the information lost when one distribution is compared to another by means of quantifying the statistical distance between two distributions [40]. Pure trait, pure metacommunity, joint metacommunity-trait and unexplained effects are calculated as proportions of total biologically relevant information (Box S1 and Box S2). Data was rarefied to smallest sample size (swamp forests; 28) and calculations bootstrapped 25 times. Results indicated no significant change compared to using all data, hence the total dataset was used for all analyses.

### Strength and direction of selection

Predictions of genus relative abundances are computed as a function of traits reflected in the CWM values and a series of constants (λ_jk_: the Lagrange Multipliers). Each multiplier quantifies the association between a unit of change for a particular trait *j* and a proportional change in predicted relative abundance *p_ik_* (the i^th^ genus in the k^th^ community) considering all other traits are constant, formally described as: 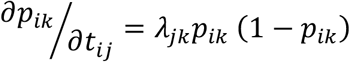 (see appendix 1 from [41]). Positive values indicate larger trait values associated with higher abundances (positive selection), negative values indicate the opposite (negative selection) with changes proportional to lambda. Values approximating zero indicate no association between specific traits and relative abundances of species. Decomposing λ_jk_ and comparing by means of a One-Way Analysis of Variance for each trait separately between forest types allows studying both the strength and direction of selection in different habitats. Note that this is done for the same constraint between forest types, as lambda values for each constraint do not scale linearly between different constraints.

### Estimation of metacommunity size

Iteratively increasing the regional species pool considered as prior in concentric circles of a fixed radius of 50 km allows estimating the spatial effect of metacommunity size. Due to computational limits, the number of permutations for the MEF calculations (see above) was reduced to two, shuffling the combinations of genera and traits at least once. Comparison of results from the analyses using all plots indicated small effects of a smaller perturbations (average of 5% difference for metacommunity effect between 5 and 50 permutations). The relationship between pure metacommunity effect and radius of metacommunity size was fitted using a smoothing loess regression (function *loess* and *predict;* R-package *stats* with span set at 0.1). Fits subsequently were used to predict values of metacommunity effect based on geographical distance to visualize general patterns for each forest type.

Exponential decay of pure metacommunity effect was described using a self-start asymptotic regression function (*SSasymp*) of the form *y(t)~y_f_+(y_0_−y_f_)e^−exp(log(α))t^* (*nls* from *stats*, [42]. A list of all packages used in R in addition to those preloaded can be found in the supplementary online material (SA2).

## Author Contributions

E.T. Pos and H. ter Steege designed the study. E.T.Pos performed analyses and took the lead in writing the manuscript, H. ter Steege supervised the writing and provided regular feedback both for the manuscript and the interpretation of the results. All other authors provided feedback on the manuscript and provided their data from the Amazon Tree Diversity Network or trait data. Authors E.T. Pos to L.V. Gamarra provided tree inventory data, authors G. Boenisch, J. Kattge, N. Kraft, A. Levesley, K. Melgaço, G. Pickavance, L. Poorter provided data on functional traits, C. Baraloto, J. Lloyd, A. A. Oliveira and H. ter Steege provided both tree inventory and functional trait data. **Competing interests**: Authors declare no competing interests. **Data and materials availability**: R scripts are available on the github repository of E.T. Pos (EdwinTPos). The data that support the findings of this study are available from The Amazon Tree Diversity Network (ATDN) upon reasonable request.

## Acknowledgements

This paper is the result of the work of hundreds of different scientists and research institutions in the Amazon over the past 80 years. Without their hard work this analysis would have been impossible. The 25-ha Long-Term Ecological Research Project of Amacayacu is a collaborative project of the Instituto Amazónico de Investigaciones Científicas Sinchi and the Universidad Nacional de Colombia Sede Medellín, in partnership with the Unidad de Manejo Especial de Parques Naturales Nacionales and the Center for Tropical Forest Science of the Smithsonian Tropical Research Institute (CTFS). The Amacayacu Forest Dynamics Plot is part of the Center for Tropical Forest Science, a global network of large-scale demographic tree plots. We acknowledge the Director and staff of the Amacayacu National Park for supporting and maintaining the project in this National Park. We would also like to thank Prof. John Harte, Prof. Bill Shipley and the anonymous reviewers for their constructive feedback during the review process.

---

**Box S1.**
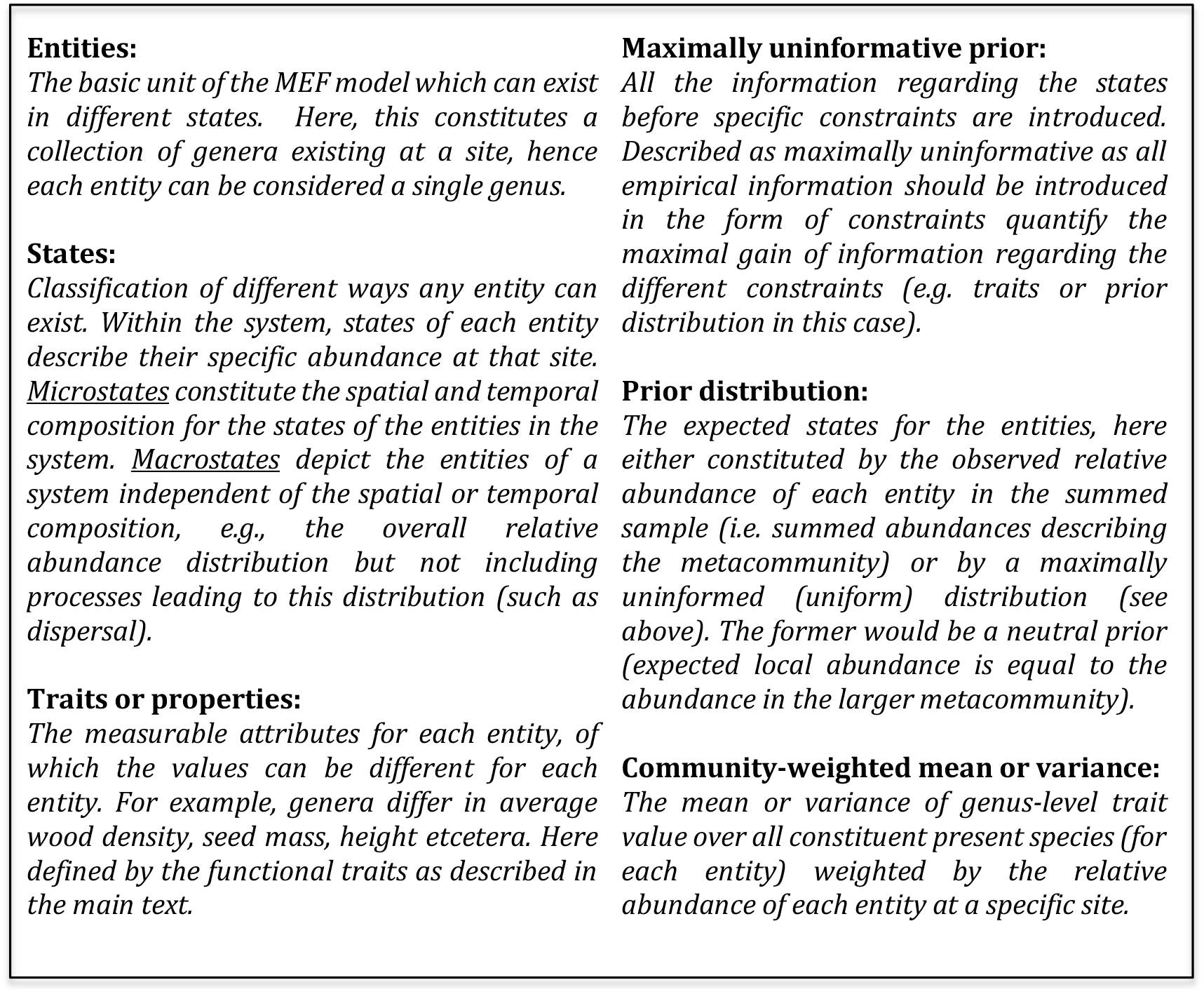
Different ingredients necessary for analyses using MEF. Definitions of the most important terms used in the MEF analyses and throughout the main text to provide the necessary framework of understanding, adapted from [1].

**Box S2.**
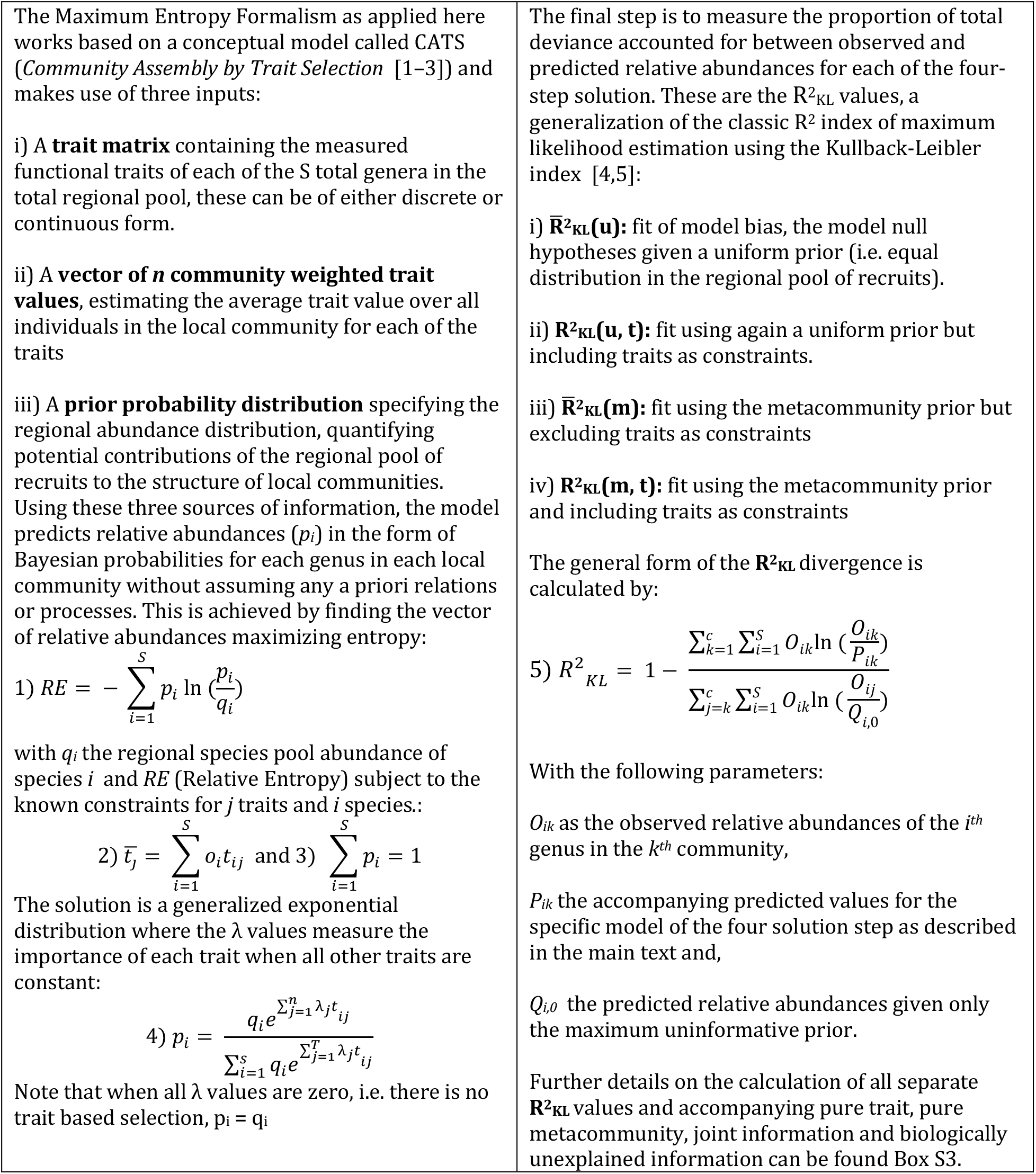
Mathematical description of the Maximum Entropy Formalism for the four-step solution. Left panel shows necessary ingredients and formulation of the Maximum Entropy Formalism. Right side panel shows decomposition of the proportion of total deviance accounted for between observed and predicted relative abundances for each of the four-step solution, adapted from [5].

**Box S3.**
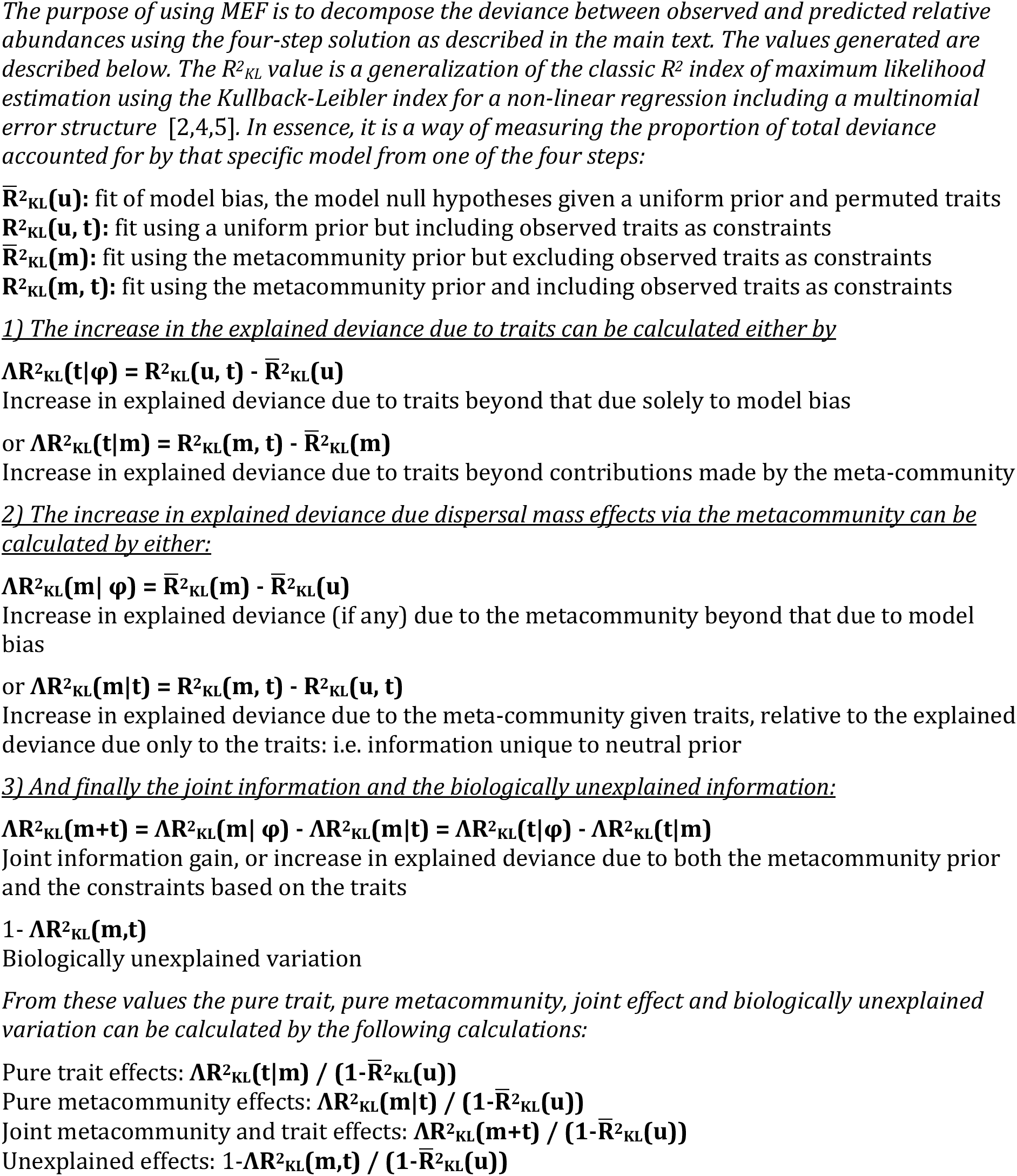
Detailed decomposition of the four-step solution from the MEF. Mathematical description of the decomposition based on the constraints and prior distributions (both uniform and neutral) for each of the steps from the four-step solution to measure the proportion of total deviance accounted for by each specific model from one of the four steps, adapted from [5].

**Fig. S1.**
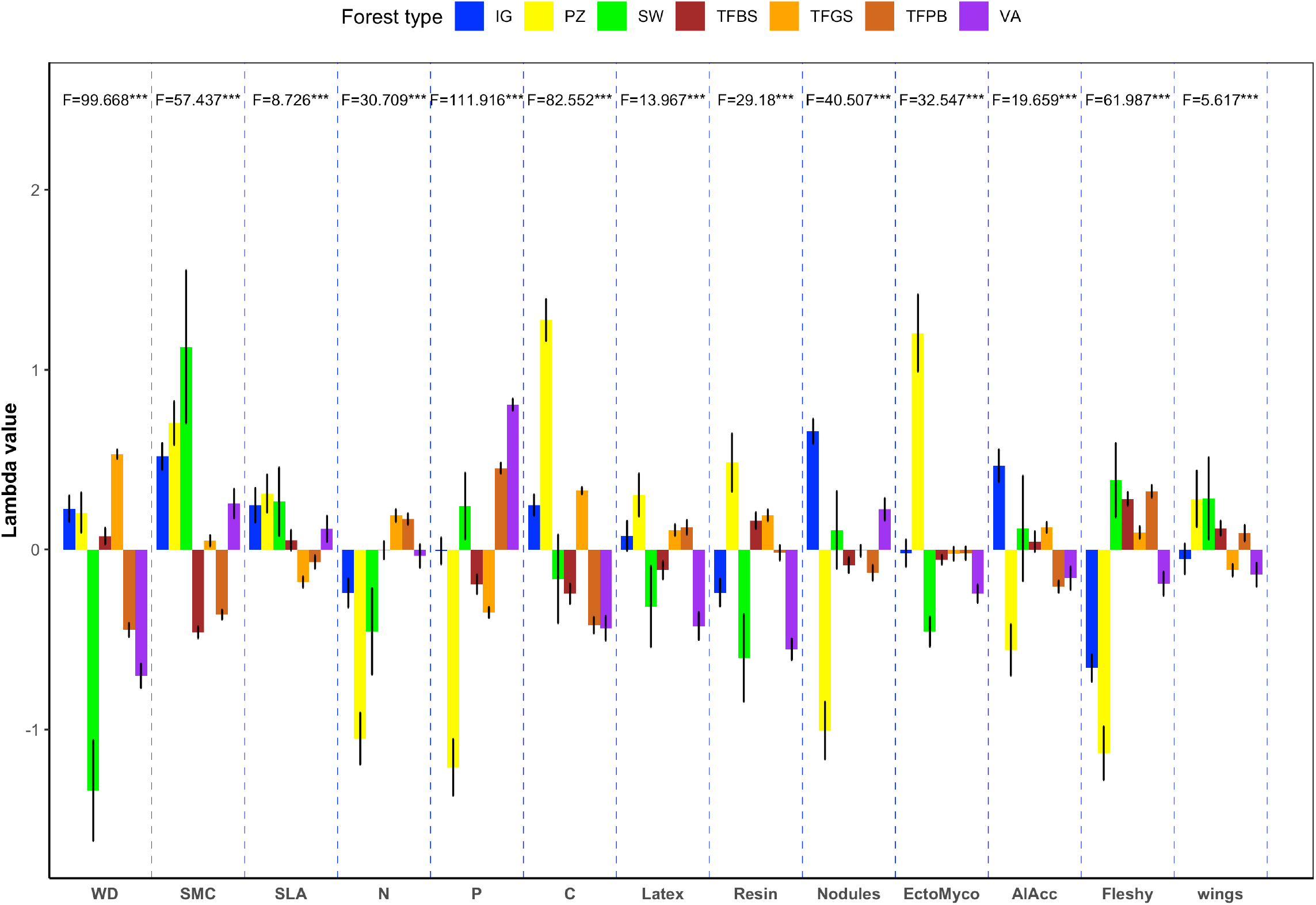
Mean lambda values with standard error bars for each functional trait and compared between forest types. Forest type abbreviations are *igapó* (IG), podzol (PZ), swamp (SW), Brazilian shield *terra firme* (TFBS), Guiana Shield *terra* firme (TFGS), Pebas *terra* firme (TFPB) and *várzea* (VA). Positive values indicate positive selection, reflective of a strong association between higher trait values and higher abundances, negative values reflect the opposite with high trait values associated with lower abundances. Differences between forest types were tested with a one-way analysis of variance with significance levels corresponding to: ns non-significant, * p < .05, ** p < .01 and *** p < .001. Abbreviations indicate functional traits: wood density (WD), seed mass class (SMC), specific leaf area (SLA), nitrogen (N), phosphorus (P) and carbon (C) leaf content. Further traits include the presence/absence of Latex, Resin, Nodules, Ectomycorrhiza (EctoMyco), the ability to accumulate aluminium (AlAcc), and the presence/absence of fleshy fruits (Fleshy) and winged seeds (Wings).

**Fig. S2.**
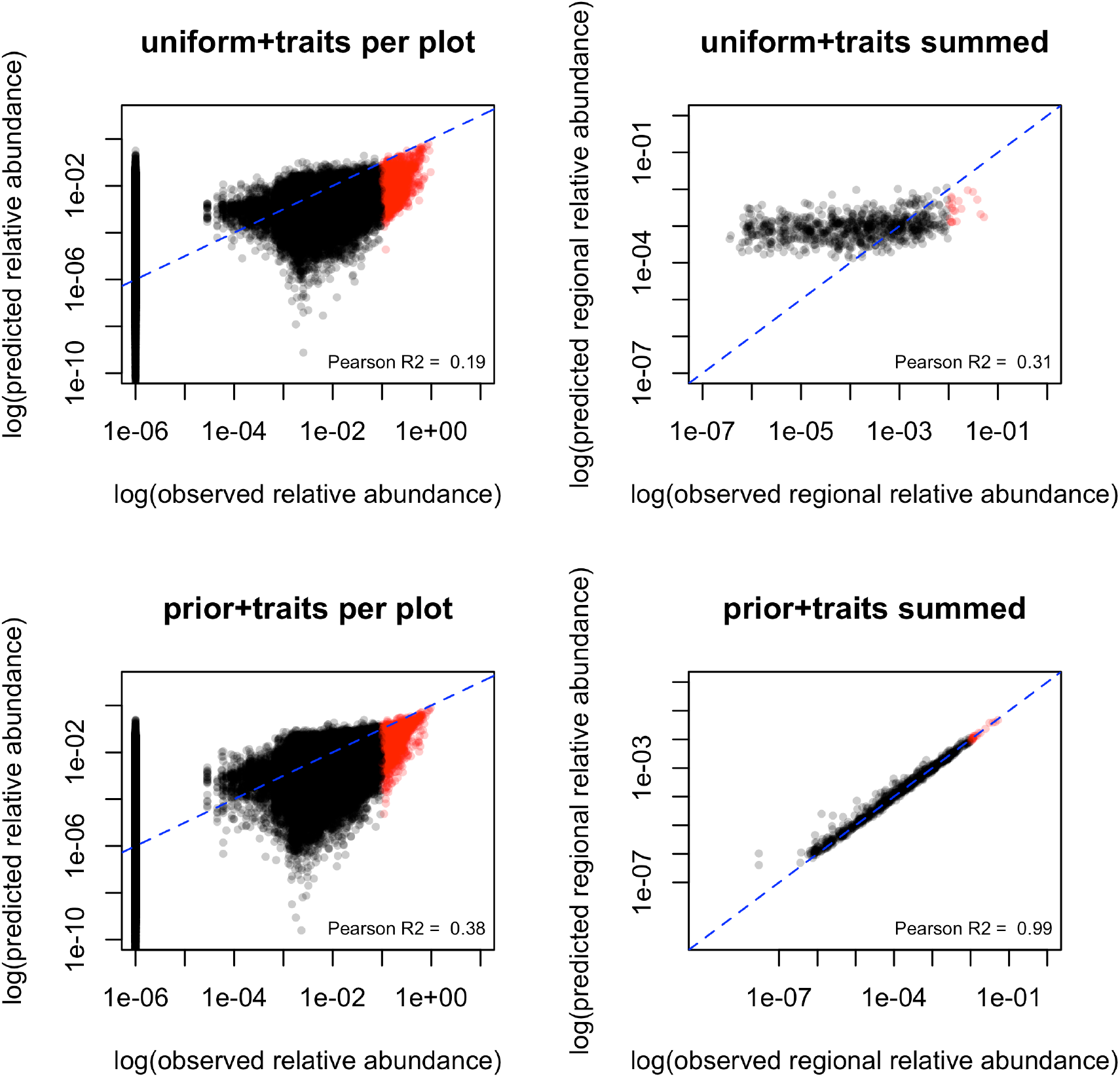
Observed relative abundances for each genus in all plots plotted against predicted relative abundance per plot (left) and summed (right) using only the traits as constraints in combination with a uniform prior (top) or the hybrid model using both traits and the metacommunity relative abundance as prior (bottom) on a log-log scale. Top figures show predictions using only a uniform prior, left separate for all plots and right for all genera summed over all plots. Bottom figures show predictions using the regional prior, again separate for all plots and genera (left) and summed over all plots for each genus (right). Red points indicate taxa with observed relative abundances over 1e-1. Lines show the x=y prediction and R^2^ values correspond to the Pearson’s correlation coefficient.

**Fig. S3.**
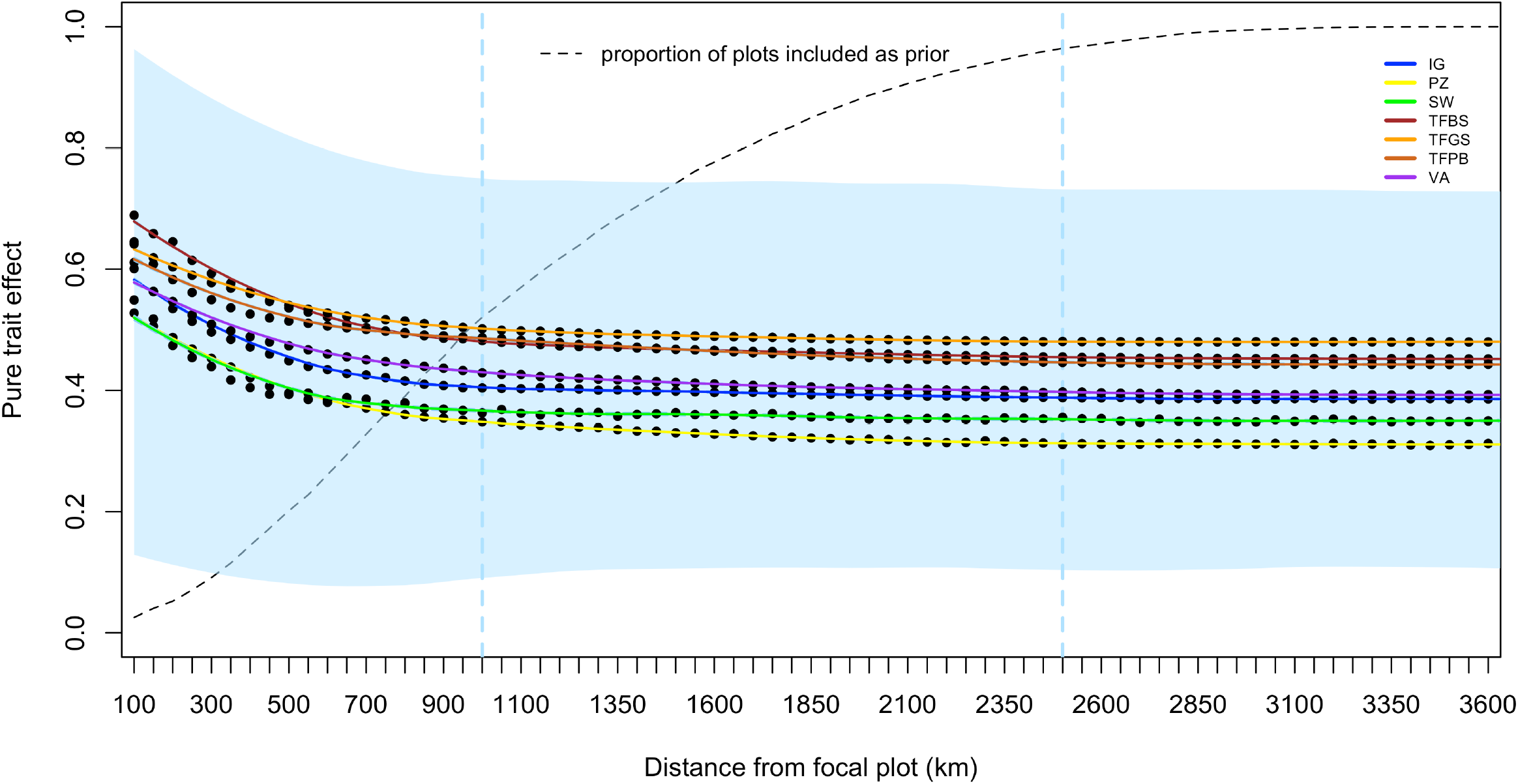
Distance decay of pure metacommunity effect. X-axis represents radius of metacommunity prior; i.e. first 100 km consists of just a few plots and at 3800 km all plots are taken into account with dashed line indicating the mean number of plots for that distance included as metacommunity prior. Y-axis represents the pure metacommunity effect, i.e. the increase in explained deviance due to the metacommunity (given traits), but relative to the explained deviance due only to the traits. It is the information unique to neutral prior taken relative to the model bias. Solid lines indicate predictions from loess regression based on all points with different colors indicating the forest types with abbreviations as in main text. Blue vertical lines indicate 1000 and 2500 km boundary points. Blue shading reflects minimum and maximum loess regression predicted values.

**Fig. S4.**
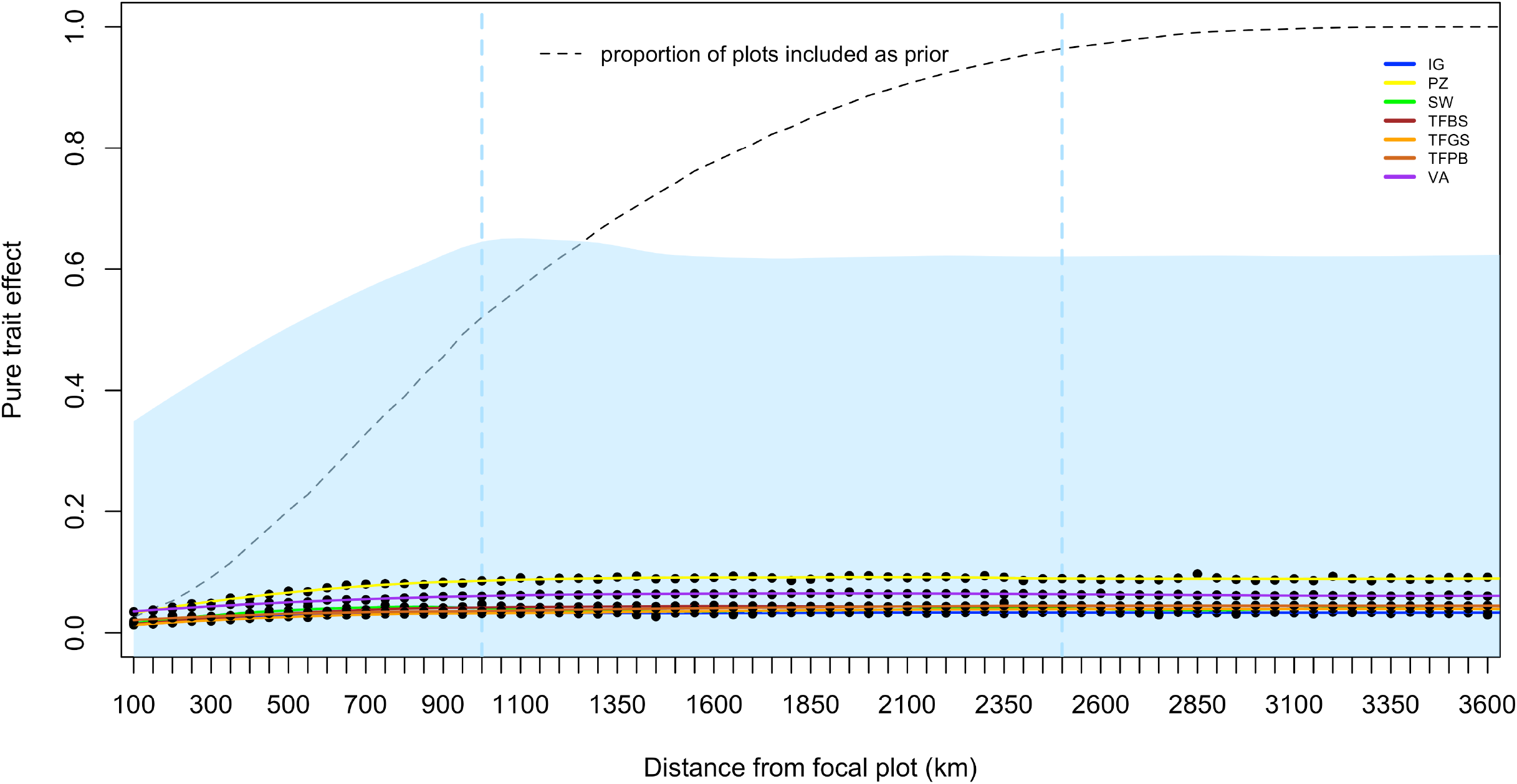
Distance decay of pure trait effect for each forest type separately and the overall dataset. X-axis represents the radius of the metacommunity prior; i.e. the first 100 km consists of just a few plots and at 3800 km all plots are taken into account. Y-axis represent the pure trait effect, i.e. the increase in explained deviance due to traits beyond contributions made by the meta-community and relative to the model bias (see also Box S2). Colors indicate the different forest types with abbreviations as in main text. Lines indicate the predictions following from the loess regression based on all points. Blue vertical lines indicate the 1000 and 2500 km boundary points. Blue shading reflects maximum values for that distance of the whole dataset.

**Fig. S5.**
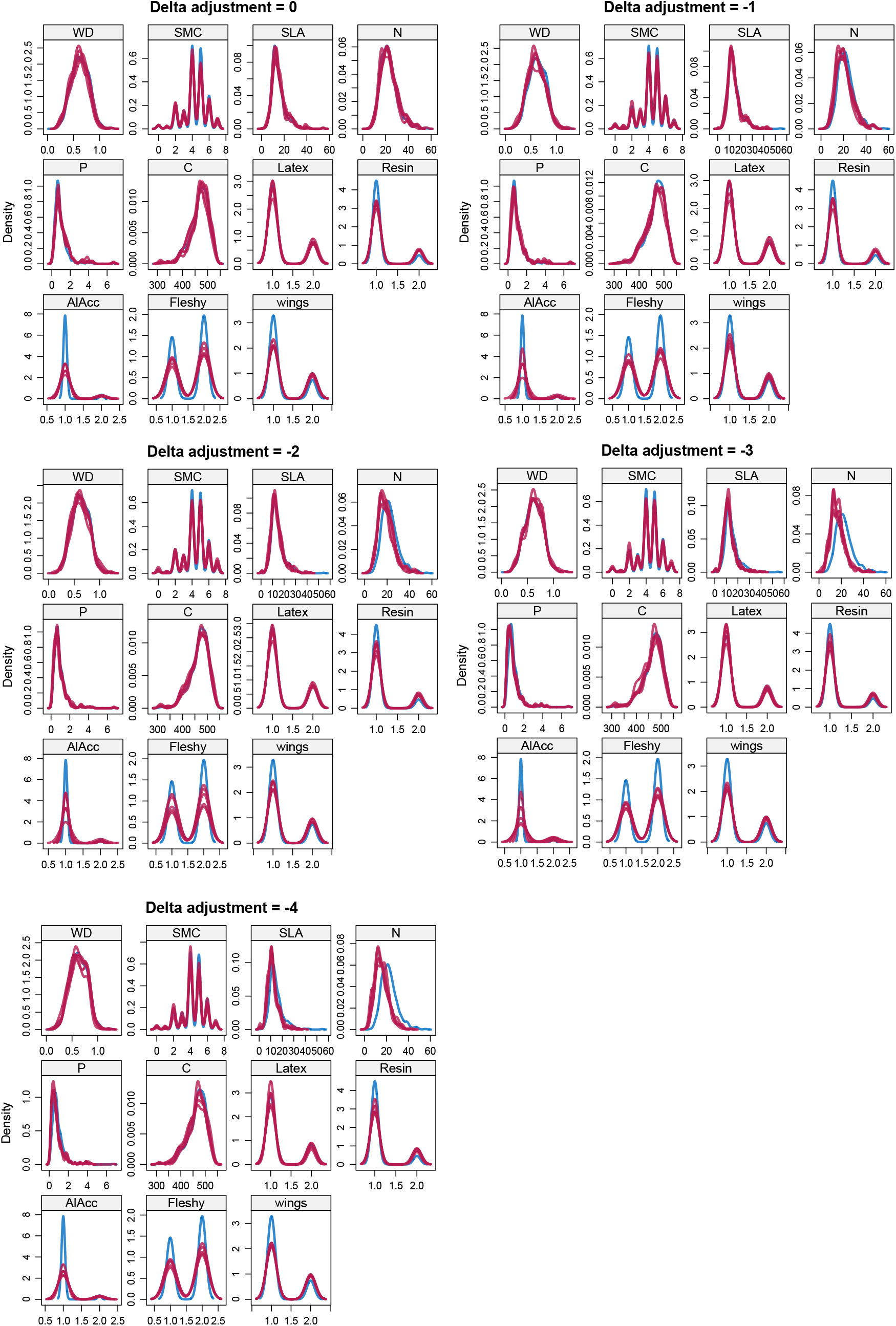
Density plots for each functional trait showing density curves for observed (blue) versus imputed data (red; 15 iterations) under the different delta adjustment scenarios. Only Nitrogen showed substantial deviation for the larger delta adjustment.

**Fig. S6.**
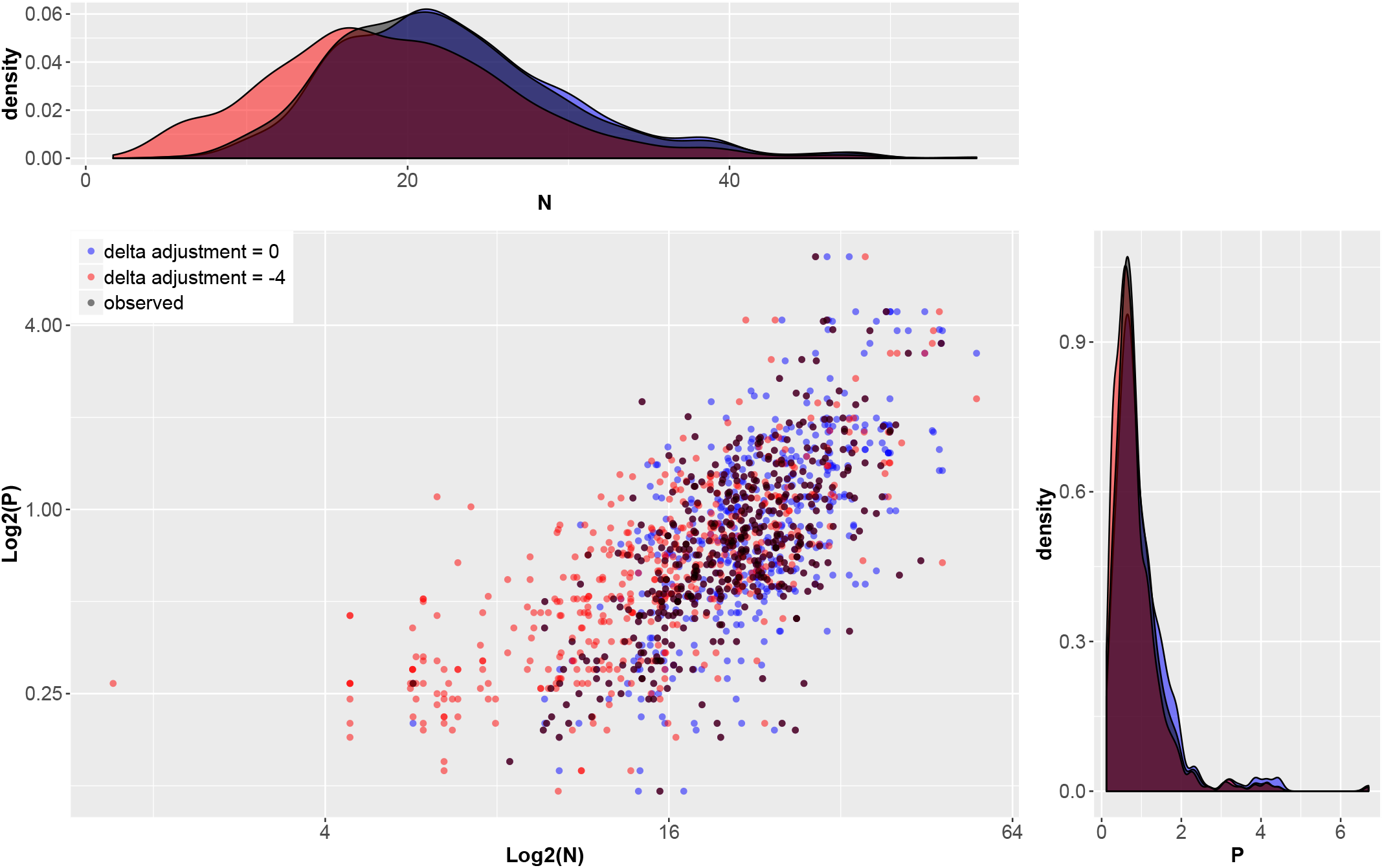
Scatterplot of observed vs. imputed leaf nitrogen and phosphorus content under different delta adjustment scenarios. As Leaf Nitrogen content showed substantial deviation in the larger delta adjustment scenario it was here plotted versus Leaf Phosphorus content, which showed hardly any deviation.

**Fig. S7.**
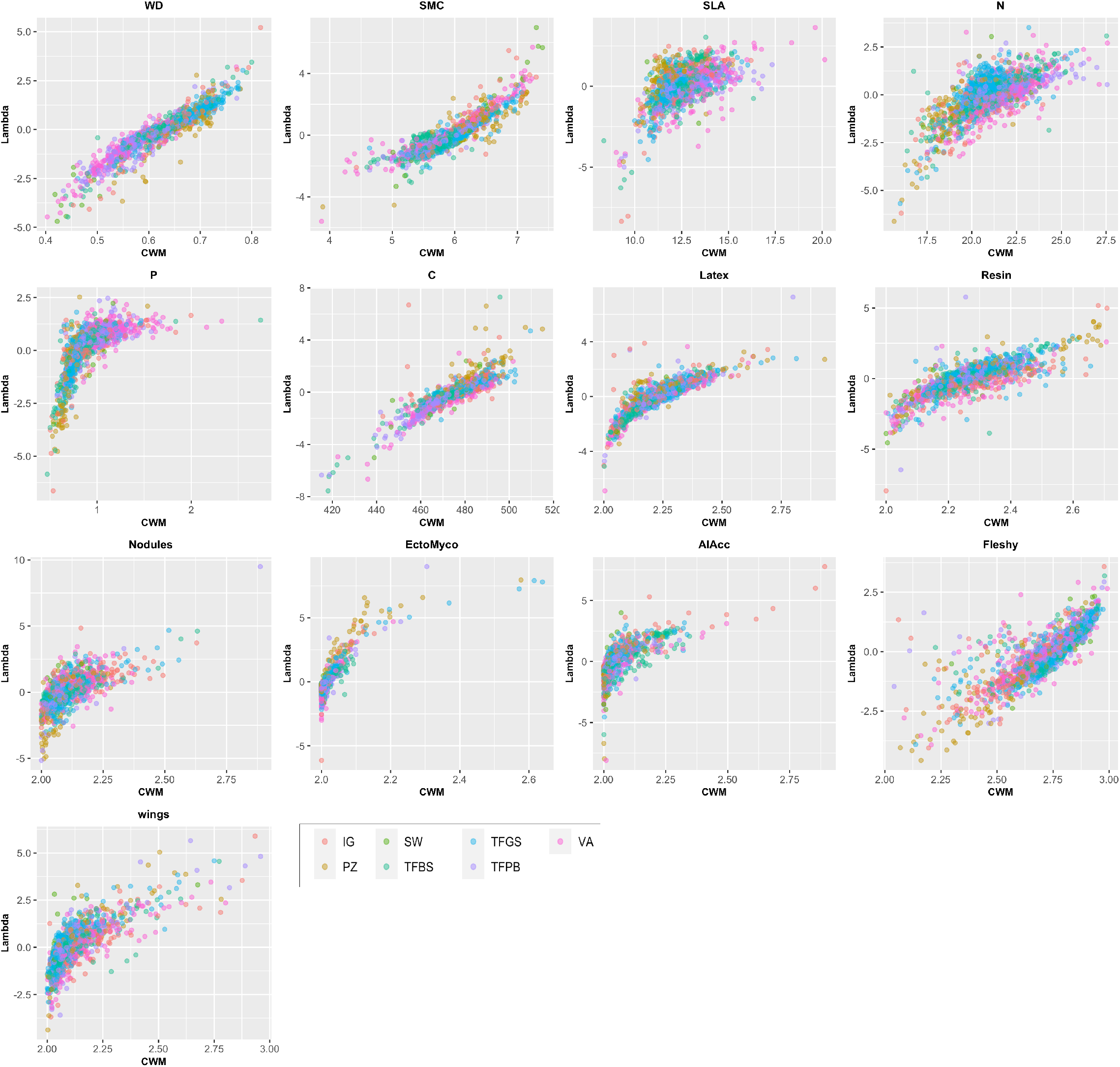
Scatterplots of the CWM values against lambda values, colored according to forest type. Titles are abbreviations for functional traits as used in the main text. Plots show in some cases these are clearly correlated (e.g. wood density, C) but for many others not (e.g. SMC and AlAcc).

**Fig. S8.**
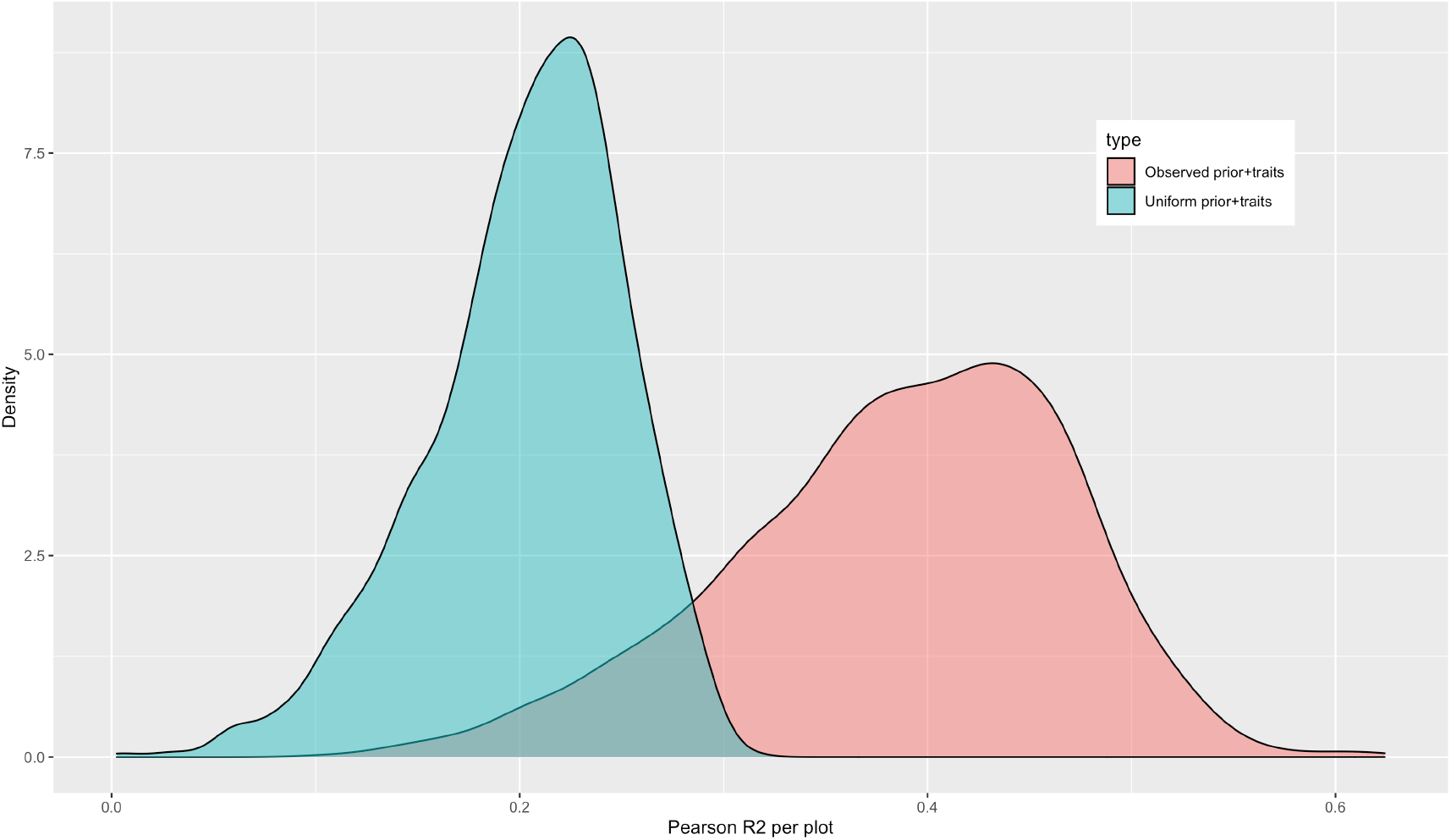
Density plot of the per-plot-Pearson correlation coefficient between predicted relative abundances of each genus. Models either used a uniform prior and functional traits (blue) or the actual observed prior and functional traits (red), results show a large increase in accuracy for the latter.

**Table S1.**
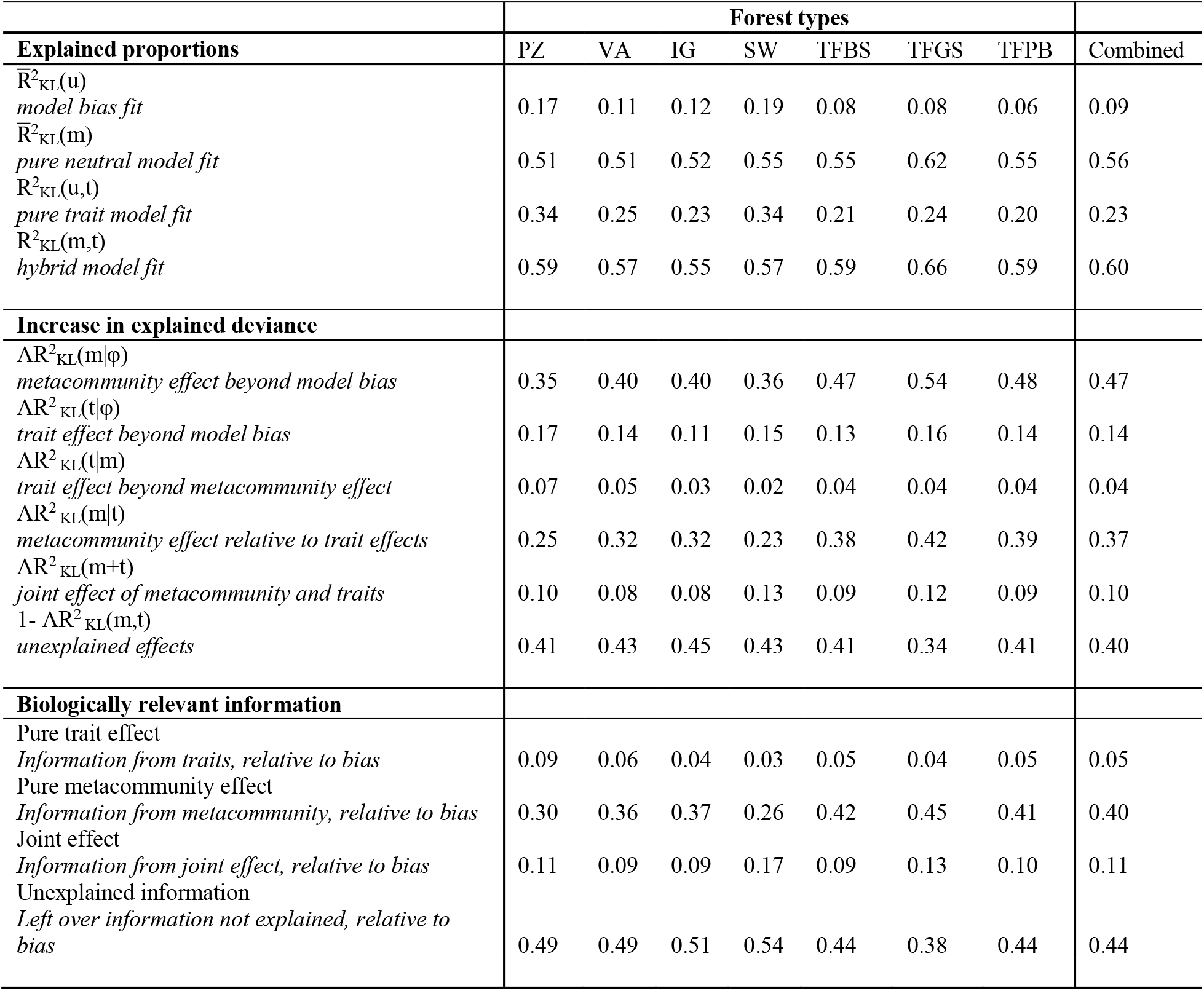
Decomposition of results from the various maximum entropy models, combined and separated by forest type (PZ podzol, IG i*gapó*, VA *várzea*, SW Swamp, TF *terra firme* with subregions BS (Brazilian Shield), GS (Guyana Shield) and PB (Pebas formation). Top rows indicate estimated proportions (R^2^_KL_) of total information reflective of variation in local relative abundance explained for by the various models. Middle rows indicate the specific information gain from any one of the used models relative to the model bias. Bottom rows show the actual effects of traits, the metacommunity and the joint information relative to the model bias.

**Table S2.** Summary statistics overview for the linear models of the various scenarios under the delta adjustment technique as described in the main text. Rows indicate the different delta adjustments used with the columns representing the standard summary statistics of the linear model comparing the imputed versus observed trait values. Results showed similar patterns with each imputation scenario, indicating a robust imputation procedure.

| Scenario | Summary statistics linear model $lm(imputed \sim observed)$ | | | | | | |
| --- | --- | --- | --- | --- | --- | --- | --- |
|  | Intercept | Std. error | T value | Pr. | R<sup>2</sup> | Adj. R<sup>2</sup> | Sigma |
| *Delta 0* | -.33 | .07 | -4.85 | 1.42e-06 | .32 | .33 | .63 |
| *Delta -2.5* | -.34 | .06 | -5.93 | 4.31e-09 | .37 | .37 | .58 |
| *Delta -5* | -.16 | .04 | -3.57 | 3.77e-04 | .40 | .40 | .53 |
| *Delta -7.5* | .64 | .09 | -7.33 | 5.18e-13 | .42 | .42 | .88 |
| *Delta -10* | 0.09 | .04 | 3.17 | 1.57e-03 | .47 | .47 | .48 |

## S-A Ecological interpretation of the MEF results

Signals of quantitative environmental selection were found to be highest for podzol forests, whereas its counterpart in the form of the dispersal mass effect from the regional pool of genera had the second lowest value. Podzol forests, having extremely nutrient poor soils could reflect a much stronger selective environment than any of the other forest types. *Terra firme* forests, presumably reflective of a less strong selective environment in terms of resource availability, showed the opposite, with approximately half of the pure trait effect in comparison with podzol forests (even when rarefied to accommodate for different sample sizes). Traits associated with protection against herbivores such as latex [7] and high leaf carbon content showed higher values associated with greater abundance on podzol soils, whereas traits indicative of investment in growth and photosynthetic ability such as high foliar concentrations of P and N [8] showed strong negative associations on nutrient poor soils. The ability to accumulate aluminium was also strongly positively associated with relative abundance on *igapó* forests, which can potentially be richer in aluminium. Lambda values also showed strong negative lambda values for wood density in swamp and forests, fitting high tree mortality and many individuals belonging to pioneer species in especially the western Amazonian swamp forests.

*Várzea* and Pebas *terra firme* forests showed a similar response. As the Pebas consists mainly of Andean sediments it has higher nutrient content, promoting lower wood density, supported by our results whereas *várzea* forests are also often flooded. There were also traits that showed no specific (strong) signal of selection on certain forest types (either positive or negative), such as latex on *igapó* and ectomycorrhiza on terra firme forests (see Fig. S1 for all lambda values). Interestingly, terra firme forests in general showed the smallest lambda values overall (positive or negative). This may be indicative of either more pronounced demographic stochasticity or ecological drift eliminating the association between traits and relative abundance. Lower effects of selection in general or more (random) variation due to the larger species pool in comparison with other forest types, however, could also be the result of mixing heterogeneous microenvironments into a single environmental class. Support for such heterogeneity within terra firme forests having influence on distribution of functional traits on valleys or plateaus has recently been found [9]. In addition, natural but also anthropogenic [10] disturbance history affects biotic community composition and can lead to changes in tree community through time, blurring relationships between traits and relative abundances. It should further be noted that, although for terra firme forests we were able to make a distinction by subregion, true within forest type heterogeneity was not taken into account. This might cause an underestimation of the deterministic effect but as of yet cannot be corrected for on this scale and is worth to be investigated in future studies. In addition, podzol forests have a smaller connected surface area and accompanying smaller number of genera in comparison with terra firme forests, adding to the calculated stronger trait effects [11,12]. When more detailed understanding and knowledge of these functional traits would be provided, this would most likely increase the explanatory power of the MEF. The fact, however, that we do not have a very specific knowledge of these interactions and specific traits is precisely the reason why the MEF can provide additional insight.

It should be noted that for species level analyses any micro environmental gradients might prove to also show (stronger) selection at local scales [13,14], as it has been shown that most variation in community composition, due to selection in regard to habitat filtering and niche conservatism, is found at lower taxonomic levels, such as between species within genera [15,16]. In contrast, theoretically it has been shown and tested that immigration numbers are actually very robust across taxonomic scales [17], validating our results of the metacommunity importance using genus level taxonomy. Spatial patterns of metacommunity effects showed shallowest declines in the centre, supporting the suggestion that high diversity of the Amazonian interior could be explained by influx of recruits due to large (overlapping) ranges. This mid-domain effect [18], however, would also predict lower species richness for the edges due to lower range overlap, assuming a closed community. This is not the case, as there is a strong species richness gradient from West (rich) to Eastern Amazonian forests (poor) [19]. The lower metacommunity effect for the edges then is most likely not due to less absolute influx of genera, but rather less influx from the Amazonian tree community. Influx from the species-rich Andes could account for the high diversity [20], yet low Amazonian metacommunity effect for Western Amazonian forests. In contrast, South-eastern parts of Amazonia receive influx from tree species-poor biomes (i.e. the Cerrado) resulting in lower diversity but also low metacommunity effect for Amazonian trees in this region.

## S-A2 List of packages used in addition to standard R [1] preloaded packages

*Vegan* Jari Oksanen, F. Guillaume Blanchet, Michael Friendly, Roeland Kindt, Pierre Legendre, Dan McGlinn, Peter R. Minchin, R. B. O’Hara, Gavin L. Simpson, Peter Solymos, M. Henry H. Stevens, Eduard Szoecs and Helene Wagner (2020). vegan: Community Ecology Package. R package version 2.5-7. https://CRAN.R-project.org/package=vegan

*FD* Laliberté, E., and P. Legendre (2010) A distance-based framework for measuring functional diversity from multiple traits. Ecology 91:299-305.

*FD* Laliberté, E., Legendre, P., and B. Shipley. (2014). FD: measuring functional diversity from multiple traits, and other tools for functional ecology. R package version 1.0-12.

*Fields* Douglas Nychka, Reinhard Furrer, John Paige, Stephan Sain (2021). “fields: Tools for spatial data.” R package version 13.3, https://github.com/dnychka/fieldsRPackage

*Binr* Sergei Izrailev (2015). binr: Cut Numeric Values into Evenly Distributed Groups. R package version 1.1. https://CRAN.R-project.org/package=binr

*doParallel* Microsoft Corporation and Steve Weston (2020). doParallel: Foreach Parallel Adaptor for the ’parallel’ Package. R package version 1.0.16. https://CRAN.R-project.org/package=doParallel

*Itertools* Steve Weston and Hadley Wickham (2014). itertools: Iterator Tools. R package version 0.1-3. https://CRAN.R-project.org/package=itertools

*Reshape* Hadley Wickham (2007). Reshaping Data with the reshape Package. Journal of Statistical Software, 21(12), 1-20. URL http://www.jstatsoft.org/v21/i12/.

*Ggplot2* H. Wickham. ggplot2: Elegant Graphics for Data Analysis. Springer-Verlag New York, 2016.

*Plotrix* Lemon, J. (2006) Plotrix: a package in the red light district of R. R-News, 6(4): 8-12.

*Mice* Stef van Buuren, Karin Groothuis-Oudshoorn (2011). mice: Multivariate Imputation by Chained Equations in R. Journal of Statistical Software, 45(3), 1-67. DOI 10.18637/jss.v045.i03.

## References and notes

[1] R. A. Chisholm and S. W. Pacala, Niche and Neutral Models Predict Asymptotically Equivalent Species Abundance Distributions in High-Diversity Ecological Communities., Proc. Natl. Acad. Sci. U. S. A. 107, 15821 (2010).

[2] D. Tilman, Niche Tradeoffs, Neutrality, and Community Structure: A Stochastic Theory of Resource Competition, Invasion, and Community Assembly., Proc. Natl. Acad. Sci. U. S. A. 101, 10854 (2004).

[3] J. Harte, Maximum Entropy and Ecology: A Theory of Abundance, Distribution, and Energetics (OUP Oxford, 2011).

[4] The Amazon Tree Diversity Network, http://atdn.myspecies.info/.

[5] S. Díaz, J. Kattge, J. H. C. Cornelissen, I. J. Wright, S. Lavorel, S. Dray, B. Reu, M. Kleyer, C. Wirth, I. Colin Prentice, E. Garnier, G. Bönisch, M. Westoby, H. Poorter, P. B. Reich, A. T. Moles, J. Dickie, A. N. Gillison, A. E. Zanne, J. Chave, S. Joseph Wright, S. N. Sheremet Ev, H. Jactel, C. Baraloto, B. Cerabolini, S. Pierce, B. Shipley, D. Kirkup, F. Casanoves, J. S. Joswig, A. Günther, V. Falczuk, N. Rüger, M. D. Mahecha, and L. D. Gorné, The Global Spectrum of Plant Form and Function, Nature 529, 167 (2016).

[6] E. T. Jaynes, Information Theory and Statistical Mechanics I, Phys. Rev. 106, 620 (1957).

[7] E. T. Jaynes, Information Theory and Statistical Mechanics. II, Phys. Rev. 108, 171 (1957).

[8] C. E. Shannon, A Mathematical Theory of Communication, Bell Syst. Tech. J. 27, 379 (1948).

[9] B. Shipley, Measuring and Interpreting Trait-Based Selection versus Meta-Community Effects during Local Community Assembly, J. Veg. Sci. 25, 55 (2014).

[10] B. Shipley, C. E. T. Paine, and C. Baraloto, Quantifying the Importance of Local Niche-Based and Stochastic Processes to Tropical Tree Community Assembly, Ecology 93, 760 (2012).

[11] J. Loranger, F. Munoz, B. Shipley, and C. Violle, What Makes Trait– Abundance Relationships When Both Environmental Filtering and Stochastic Neutral Dynamics Are at Play?, Oikos 127, 1735 (2018).

[12] K. G. Dexter, M. Lavin, B. M. Torke, A. D. Twyford, T. A. Kursar, P. D. Coley, C. Drake, R. Hollands, and R. T. Pennington, Dispersal Assembly of Rain Forest Tree Communities across the Amazon Basin, Proc. Natl. Acad. Sci. 114, 2645 (2017).

[13] H. ter Steege, N. C. A. Pitman, D. Sabatier, C. Baraloto, R. P. Salomão, J. E. Guevara, O. L. Phillips, C. V Castilho, W. E. Magnusson, J.-F. Molino, A. Monteagudo, P. Núñez Vargas, J. C. Montero, T. R. Feldpausch, E. N. H. Coronado, T. J. Killeen, B. Mostacedo, R. Vasquez, R. L. Assis, J. Terborgh, F. Wittmann, A. Andrade, W. F. Laurance, S. G. W. Laurance, B. S. Marimon, B.-H. Marimon, I. C. Guimarães Vieira, I. L. Amaral, R. Brienen, H. Castellanos, D. Cárdenas López, J. F. Duivenvoorden, H. F. Mogollón, F. D. de A. Matos, N. Dávila, R. García-Villacorta, P. R. Stevenson Diaz, F. Costa, T. Emilio, C. Levis, J. Schietti, P. Souza, A. Alonso, F. Dallmeier, A. J. D. Montoya, M. T. Fernandez Piedade, A. Araujo-Murakami, L. Arroyo, R. Gribel, P. V. A. Fine, C. A. Peres, M. Toledo, G. A. Aymard C, T. R. Baker, C. Cerón, J. Engel, T. W. Henkel, P. Maas, P. Petronelli, J. Stropp, C. E. Zartman, D. Daly, D. Neill, M. Silveira, M. R. Paredes, J. Chave, D. de A. Lima Filho, P. M. Jørgensen, A. Fuentes, J. Schöngart, F. Cornejo Valverde, A. Di Fiore, E. M. Jimenez, M. C. Peñuela Mora, J. F. Phillips, G. Rivas, T. R. van Andel, P. von Hildebrand, B. Hoffman, E. L. Zent, Y. Malhi, A. Prieto, A. Rudas, A. R. Ruschell, N. Silva, V. Vos, S. Zent, A. A. Oliveira, A. C. Schutz, T. Gonzales, M. Trindade Nascimento, H. Ramirez-Angulo, R. Sierra, M. Tirado, M. N. Umaña Medina, G. van der Heijden, C. I. A. Vela, E. Vilanova Torre, C. Vriesendorp, O. Wang, K. R. Young, C. Baider, H. Balslev, C. Ferreira, I. Mesones, A. Torres-Lezama, L. E. Urrego Giraldo, R. Zagt, M. N. Alexiades, L. Hernandez, I. Huamantupa-Chuquimaco, W. Milliken, W. Palacios Cuenca, D. Pauletto, E. Valderrama Sandoval, L. Valenzuela Gamarra, K. G. Dexter, K. Feeley, G. Lopez-Gonzalez, M. R. Silman, S. P. Hubbell, F. He, R. Condit, L. Borda-de-Agua, J. Kellner, H. Ter Steege, G. A. Black, T. H. Dobzhansky, C. Pavan, J. M. Pires, T. Dobzhansky, G. A. Black, M. J. G. Hopkins, M. J. Costello, R. M. May, N. E. Stork, P. Haripersaud, H. ter Steege, J.-J. de Granville, H. Chevillotte, M. Hoff, D. P. Bebber, M. A. Carine, J. R. Wood, A. H. Wortley, D. J. Harris, G. T. Prance, G. Davidse, J. Paige, T. D. Pennington, N. K. Robson, R. W. Scotland, B. J. McGill, R. S. Etienne, J. S. Gray, D. Alonso, M. J. Anderson, H. K. Benecha, M. Dornelas, B. J. Enquist, J. L. Green, F. He, A. H. Hurlbert, A. E. Magurran, P. A. Marquet, B. A. Maurer, A. Ostling, C. U. Soykan, K. I. Ugland, E. P. White, R. J. Warren, D. K. Skelly, O. J. Schmitz, M. A. Bradford, N. C. A. Pitman, J. W. Terborgh, M. R. Silman, P. N. V, D. A. Neill, C. E. Cerón, W. A. Palacios, M. Aulestia, N. C. A. Pitman, M. R. Silman, J. W. Terborgh, F. D. Lozano, M. W. Schwartz, M. W. Schwartz, D. Simberloff, J. E. Richardson, R. T. Pennington, T. D. Pennington, P. M. Hollingsworth, T. L. Couvreur, F. Forest, W. J. Baker, S. Cavers, C. W. Dick, D. H. Janzen, S. A. Mangan, S. A. Schnitzer, E. A. Herre, K. M. Mack, M. C. Valencia, E. I. Sanchez, J. D. Bever, W. Balée, D. G. Campbell, C. Levis, P. F. de Souza, J. Schietti, T. Emilio, J. L. P. V. Pinto, C. R. Clement, F. R. C. Costa, D. A. Posey, E. Montoya, V. Rull, N. D. Stansell, M. B. Abbott, S. Nogué, B. W. Bird, W. A. Díaz, C. Gomez-Navarro, C. Jaramillo, F. Herrera, S. L. Wing, R. Callejas, C. H. McMichael, D. R. Piperno, M. B. Bush, M. R. Silman, A. R. Zimmerman, M. F. Raczka, L. C. Lobato, H. ter Steege, P. P. Haripersaud, O. S. Bánki, F. Schieving, S. J. Phillips, R. P. Anderson, R. E. Schapire, S. J. Phillips, M. Dudik, C. A. Quesada, J. Lloyd, L. O. Anderson, N. M. Fyllas, M. Schwarz, C. I. Czimczik, B. Rollet, H. ter Steege, N. C. Pitman, O. L. Phillips, J. Chave, D. Sabatier, A. Duque, J. F. Molino, M. F. Prévost, R. Spichiger, H. Castellanos, P. von Hildebrand, R. Vásquez, P. M. Fearnside, D. Mouillot, D. R. Bellwood, C. Baraloto, J. Chave, R. Galzin, M. Harmelin-Vivien, M. Kulbicki, S. Lavergne, S. Lavorel, N. Mouquet, C. E. Paine, J. Renaud, W. Thuiller, G. Lopez-Gonzalez, S. L. Lewis, M. Burkitt, O. L. Phillips, P. J. M. Maas, L. Y. T. Westra, H. Rainer, A. Q. Lobão, R. H. J. Erkens, P. V. A. Fine, D. C. Daly, G. V. Muñoz, I. Mesones, K. M. Cameron, K. J. Feeley, M. R. Silman, M. Dufrene, P. Legendre, J. Bunge, M. Fitzpatrick, J. Bunge, L. Woodard, D. Böhning, J. A. Foster, S. Connolly, H. K. Allen, J.-P. Z. Wang, B. G. Lindsay, A. Chao, R. K. Colwell, C.-W. Lin, N. J. Gotelli, C. X. Mao, R. K. Colwell, U. Brose, N. D. Martinez, R. J. Williams, I. J. Good, A. Chao, S.-M. Lee, I. Rocchetti, J. Bunge, D. Bohning, R. A. Fisher, A. S. Corbet, C. B. Williams, F. W. Preston, J.-P. Wang, P. V. A. Fine, and R. H. Ree, Hyperdominance in the Amazonian Tree Flora, Science 342, 1243092 (2013).

[14] H. ter Steege, P. I. Prado, R. A. F. d. Lima, E. Pos, L. de Souza Coelho, D. de Andrade Lima Filho, R. P. Salomão, I. L. Amaral, F. D. de Almeida Matos, C. V. Castilho, O. L. Phillips, J. E. Guevara, M. de Jesus Veiga Carim, D. Cárdenas López, W. E. Magnusson, F. Wittmann, M. P. Martins, D. Sabatier, M. V. Irume, J. R. da Silva Guimarães, J. F. Molino, O. S. Bánki, M. T. F. Piedade, N. C. A. Pitman, J. F. Ramos, A. Monteagudo Mendoza, E. M. Venticinque, B. G. Luize, P. Núñez Vargas, T. S. F. Silva, E. M. M. de Leão Novo, N. F. C. Reis, J. Terborgh, A. G. Manzatto, K. R. Casula, E. N. Honorio Coronado, J. C. Montero, A. Duque, F. R. C. Costa, N. Castaño Arboleda, J. Schöngart, C. E. Zartman, T. J. Killeen, B. S. Marimon, B. H. Marimon-Junior, R. Vasquez, B. Mostacedo, L. O. Demarchi, T. R. Feldpausch, J. Engel, P. Petronelli, C. Baraloto, R. L. Assis, H. Castellanos, M. F. Simon, M. B. de Medeiros, A. Quaresma, S. G. W. Laurance, L. M. Rincón, A. Andrade, T. R. Sousa, J. L. Camargo, J. Schietti, W. F. Laurance, H. L. de Queiroz, H. E. M. Nascimento, M. A. Lopes, E. de Sousa Farias, J. L. L. Magalhães, R. Brienen, G. A. Aymard C, J. D. C. Revilla, I. C. G. Vieira, B. B. L. Cintra, P. R. Stevenson, Y. O. Feitosa, J. F. Duivenvoorden, H. F. Mogollón, A. Araujo-Murakami, L. V. Ferreira, J. R. Lozada, J. A. Comiskey, J. J. de Toledo, G. Damasco, N. Dávila, A. Lopes, R. García-Villacorta, F. Draper, A. Vicentini, F. Cornejo Valverde, J. Lloyd, V. H. F. Gomes, D. Neill, A. Alonso, F. Dallmeier, F. C. de Souza, R. Gribel, L. Arroyo, F. A. Carvalho, D. P. P. de Aguiar, D. D. do Amaral, M. P. Pansonato, K. J. Feeley, E. Berenguer, P. V. A. Fine, M. C. Guedes, J. Barlow, J. Ferreira, B. Villa, M. C. Peñuela Mora, E. M. Jimenez, J. C. Licona, C. Cerón, R. Thomas, P. Maas, M. Silveira, T. W. Henkel, J. Stropp, M. R. Paredes, K. G. Dexter, D. Daly, T. R. Baker, I. Huamantupa-Chuquimaco, W. Milliken, T. Pennington, J. S. Tello, J. L. M. Pena, C. A. Peres, B. Klitgaard, A. Fuentes, M. R. Silman, A. Di Fiore, P. von Hildebrand, J. Chave, T. R. van Andel, R. R. Hilário, J. F. Phillips, G. Rivas-Torres, J. C. Noronha, A. Prieto, T. Gonzales, R. de Sá Carpanedo, G. P. G. Gonzales, R. Z. Gómez, D. de Jesus Rodrigues, E. L. Zent, A. R. Ruschel, V. A. Vos, É. Fonty, A. B. Junqueira, H. P. D. Doza, B. Hoffman, S. Zent, E. M. Barbosa, Y. Malhi, L. C. de Matos Bonates, I. P. de Andrade Miranda, N. Silva, F. R. Barbosa, C. I. A. Vela, L. F. M. Pinto, A. Rudas, B. W. Albuquerque, M. N. Umaña, Y. A. Carrero Márquez, G. van der Heijden, K. R. Young, M. Tirado, D. F. Correa, R. Sierra, J. B. P. Costa, M. Rocha, E. Vilanova Torre, O. Wang, A. A. Oliveira, M. Kalamandeen, C. Vriesendorp, H. Ramirez-Angulo, M. Holmgren, M. T. Nascimento, D. Galbraith, B. M. Flores, V. V. Scudeller, A. Cano, M. A. Ahuite Reategui, I. Mesones, C. Baider, C. Mendoza, R. Zagt, L. E. Urrego Giraldo, C. Ferreira, D. Villarroel, R. Linares-Palomino, W. Farfan-Rios, W. Farfan-Rios, L. F. Casas, S. Cárdenas, H. Balslev, A. Torres-Lezama, M. N. Alexiades, K. Garcia-Cabrera, L. Valenzuela Gamarra, E. H. Valderrama Sandoval, F. Ramirez Arevalo, L. Hernandez, A. F. Sampaio, S. Pansini, W. Palacios Cuenca, E. A. de Oliveira, D. Pauletto, A. Levesley, K. Melgaço, and G. Pickavance, Biased-Corrected Richness Estimates for the Amazonian Tree Flora, Sci. Rep. 10, 1 (2020).

[15] Tropicos, Tropicos.Org. Missouri Botanical Garden. Accessed 2018, Version Number Unknown Due to Lack of Release-History., (unpublished).

[16] B. Boyle, N. Hopkins, Z. Lu, J. A. Raygoza Garay, D. Mozzherin, T. Rees, N. Matasci, M. L. Narro, W. H. Piel, S. J. McKay, S. Lowry, C. Freeland, R. K. Peet, and B. J. Enquist, The Taxonomic Name Resolution Service: An Online Tool for Automated Standardization of Plant Names., BMC Bioinformatics 14, 16 (2013).

[17] T. Haugaasen and C. Peres, Floristic, Edaphic and Structural Characteristics of Flooded and Unflooded Forests in the Lower Rio Purús Region of Central Amazonia, Brazil, Acta Amaz. 36, 25 (2006).

[18] J. Chave, D. Coomes, and S. Jansen, Towards a Worldwide Wood Economics Spectrum, Ecol. Lett. 351 (2009).

[19] N. J. B. Kraft, R. Valencia, and D. D. Ackerly, Functional Traits and Niche-Based Tree Community Assembly in an Amazonian Forest., Science 322, 580 (2008).

[20] N. M. Fyllas, S. Patino, T. R. Baker, G. Bielefeld Nardoto, L. A. Martinelli, C. A. Quesada, R. Paiva, M. Schwarz, V. Horna, L. M. Mercado, A. Santos, L. Arroyo, E. M. Jiměnez, F. J. Luizao, D. A. Neill, N. Silva, A. Prieto, A. Rudas, M. Silviera, I. C. G. Vieira, G. Lopez-Gonzalez, Y. Malhi, O. L. Phillips, and J. Lloyd, Basin-Wide Variations in Foliar Properties of Amazonian Forest: Phylogeny, Soils and Climate, Biogeosciences 6, 2677 (2009).

[21] C. Baraloto, C. E. T. Paine, L. Poorter, J. Beauchene, D. Bonal, A. M. Domenach, B. Hérault, S. Patiño, J. C. Roggy, and J. Chave, Decoupled Leaf and Stem Economics in Rain Forest Trees, Ecol. Lett. 13, 1338 (2010).

[22] C. Fortunel, P. V. A. Fine, and C. Baraloto, Leaf, Stem and Root Tissue Strategies across 758 Neotropical Tree Species, Funct. Ecol. 26, 1153 (2012).

[23] C. E. T. Paine, C. Baraloto, J. Chave, and B. Hérault, Functional Traits of Individual Trees Reveal Ecological Constraints on Community Assembly in Tropical Rain Forests, Oikos 120, 720 (2011).

[24] S. Patiño, N. M. Fyllas, T. R. Baker, R. Paiva, C. A. Quesada, A. J. B. Santos, M. Schwarz, H. ter Steege, O. L. Phillips, and J. Lloyd., Coordination of Physiological and Structural Traits in Amazon Forest Trees., Biogeosciences 9, 775 (2012).

[25] F. M. J. Ohler, Phytomass and Mineral Content in Untouched Forest, (1980).

[26] J. Thompson, J. Proctor, V. Viana, W. Milliken, J. A. Ratter, and D. A. Scott, Ecological Studies on a Lowland Evergreen Rain Forest on Maraca Island, Roraima, Brazil. I. Physical Environment, Forest Structure and Leaf Chemistry, J. Ecol. 689 (1992).

[27] T. L. Pons, K. Perreijn, C. Van Kessel, and M. J. A. Werger, Symbiotic Nitrogen Fixation in a Tropical Rainforest: 15N Natural Abundance Measurements Supported by Experimental Isotopic Enrichment, New Phytol. 173, 154 (2006).

[28] S. Foster and C. H. Janson, The Relationship between Seed Size and Establishment Conditions in Tropical Woody Plants, Ecology 66, 773 (1985).

[29] D. S. Hammond and V. K. Brown, Hammond and Brown 1995 Seed Size. Ecology.Pdf, Ecology 76, 2544 (1995).

[30] Royal Botanic Gardens Kew, Seed Information Database (SID), (unpublished).

[31] H. ter Steege and D. Hammond, Character Convergence, Diversity, and Disturbance in Tropical Rain Forest in Guyana, Ecology 82, 3197 (2001).

[32] L. Tedersoo and M. C. Brundrett, Evolution of Ectomycorrhizal Symbiosis in Plants., Biogeogr. Mycorrhizal Symbiosis. Springer Int. Publ. Basel 407 (2017).

[33] J. I. Sprent, Nodulation in Legumes, (unpublished).

[34] P. S. Soltis, D. E. Soltis, and M. W. Chase, Angiosperm Phylogeny Inferred from Multiple Genes as a Tool for Comparative Biology., Nature 402, 402 (1999).

[35] S. Jansen, T. Watanabe, S. Dessein, E. Smets, and E. Robbrecht, A Comparative Study of Metal Levels in Leaves of Some Al-Accumulating Rubiaceae, Ann. Bot. 91, 657 (2003).

[36] S. Jansen, T. Watanabe, and E. Smets., Aluminium Accumulation in Leaves of 127 Species in Melastomataceae, with Comments on the Order Myrtales, Ann. Bot. 90, 53 (2002).

[37] G. Sonnier, B. Shipley, and M. L. Navas, Quantifying Relationships between Traits and Explicitly Measured Gradients of Stress and Disturbance in Early Successional Plant Communities, J. Veg. Sci. 21, 1014 (2010).

[38] S. van Buuren and K. Groothuis-Oudshoorn, Mice: Multivariate Imputation by Chained Equations in R, J. Stat. Softw. 45, 1 (2011).

[39] S. H. Roxburgh and K. Mokany, On Testing Predictions of Species Relative Abundance from Maximum Entropy Optimisation, Oikos 119, 583 (2010).

[40] A. Colin Cameron and F. A. G. Windmeijer, An R-Squared Measure of Goodness of Fit for Some Common Nonlinear Regression Models, J. Econom. 77, 329 (1997).

[41] G. Sonnier, M. L. Navas, A. Fayolle, and B. Shipley, Quantifying Trait Selection Driving Community Assembly: A Test in Herbaceous Plant Communities under Contrasted Land Use Regimes, Oikos 121, 1103 (2012).

[42] R Core Team, R: A Language and Environment for Statistical Computing.

[43] E. Weiher, A. van der Werf, K. Thompson, M. Roderick, E. Garnier, and O. Eriksson, Challenging Theophrastus: A Common Core List of Plant Traits for Functional Ecology, J. Veg. Sci. 10, 609 (1999).

[44] K. G. Raghothama, Phosphate Acquisition, Annu. Rev. Plant Physiol. Plant Mol. Biol. 50, 665 (1999).

[45] L. Poorter, M. Van De Plassche, S. Willems, and R. G. A. Boot, Leaf Traits and Herbivory Rates of Tropical Tree Species Differing in Successional Status, Plant Biol. 6, 746 (2004).

[46] A. A. Agrawal, Macroevolution of Plant Defense Strategies, Trends Ecol. Evol. 22, 103 (2007).

[47] J. I. Sprent and R. Parsons, Nitrogen Fixation in Legume and Non-Legume Trees, F. Crop. Res. 65, 183 (2000).

[48] J. V. Colpaert, J. H. L. Wevers, E. Krznaric, and K. Adriaensen, How Metal-Tolerant Ecotypes of Ectomycorrhizal Fungi Protect Plants from Heavy Metal Pollution, Ann. For. Sci. 68, 17 (2011).

[49] C. Poschenrieder, B. Gunsé, I. Corrales, and J. Barceló, A Glance into Aluminum Toxicity and Resistance in Plants, Sci. Total Environ. 400, 356 (2008).

[50] H. F. Howe and J. Smallwood, Ecology of Seed Dispersal Ecology of Seed Dispersal, 13, 201 (2013).

## REFERENCES SUPPLEMENTARY MATERIAL

[1] B. Shipley, From Plant Traits to Vegetation Structure. Chance and Selection in the Assembly of Ecological Communities (Cambridge University Press, 2010).

[2] B. Shipley, C. E. T. Paine, and C. Baraloto, Quantifying the Importance of Local Niche-Based and Stochastic Processes to Tropical Tree Community Assembly, Ecology 93, 760 (2012).

[3] J. Loranger, F. Munoz, B. Shipley, and C. Violle, What Makes Trait– Abundance Relationships When Both Environmental Filtering and Stochastic Neutral Dynamics Are at Play?, Oikos 127, 1735 (2018).

[4] A. Colin Cameron and F. A. G. Windmeijer, An R-Squared Measure of Goodness of Fit for Some Common Nonlinear Regression Models, J. Econom. 77, 329 (1997).

[5] B. Shipley, Measuring and Interpreting Trait-Based Selection versus Meta-Community Effects during Local Community Assembly, J. Veg. Sci. 25, 55 (2014).

[6] H. ter Steege, N. C. a Pitman, O. L. Phillips, J. Chave, D. Sabatier, A. Duque, J.-F. Molino, M.-F. Prévost, R. Spichiger, H. Castellanos, P. von Hildebrand, and R. Vásquez, Continental-Scale Patterns of Canopy Tree Composition and Function across Amazonia., Nature 443, 444 (2006).

[7] A. A. Agrawal, Macroevolution of Plant Defense Strategies, Trends Ecol. Evol. 22, 103 (2007).

[8] J. C. Ordoñez, P. M. Van Bodegom, J. P. M. Witte, I. J. Wright, P. B. Reich, and R. Aerts, A Global Study of Relationships between Leaf Traits, Climate and Soil Measures of Nutrient Fertility, Glob. Ecol. Biogeogr. 18, 137 (2009).

[9] R. S. Oliveira, F. R. C. Costa, E. van Baalen, A. de Jonge, P. R. Bittencourt, Y. Almanza, F. de V. Barros, E. C. Cordoba, M. V. Fagundes, S. Garcia, Z. T. T. M. Guimaraes, M. Hertel, J. Schietti, J. Rodrigues-Souza, and L. Poorter, Embolism Resistance Drives the Distribution of Amazonian Rainforest Tree Species along Hydro-Topographic Gradients, New Phytol. (2018).

[10] C. Levis, B. M. Flores, P. A. Moreira, B. G. Luize, R. P. Alves, J. Franco-Moraes, J. Lins, E. Konings, M. Peña-Claros, F. Bongers, F. R. C. Costa, and C. R. Clement, How People Domesticated Amazonian Forests, Front. Ecol. Evol. 5, (2018).

[11] J. E. Guevara, G. Damasco, C. Baraloto, P. V. A. Fine, M. C. Peñuela, C. Castilho, A. Vincentini, D. Cárdenas, F. Wittmann, N. Targhetta, O. Phillips, J. Stropp, I. Amaral, P. Maas, A. Monteagudo, E. M. Jimenez, R. Thomas, R. Brienen, Á. Duque, W. Magnusson, C. Ferreira, E. Honorio, F. de Almeida Matos, F. R. Arevalo, J. Engel, P. Petronelli, R. Vasquez, and H. ter Steege, Low Phylogenetic Beta Diversity and Geographic Neo-Endemism in Amazonian White-Sand Forests, Biotropica 48, 34 (2016).

[12] H. ter Steege, N. C. A. Pitman, D. Sabatier, C. Baraloto, R. P. Salomão, J. E. Guevara, O. L. Phillips, C. V Castilho, W. E. Magnusson, J.-F. Molino, A. Monteagudo, P. Núñez Vargas, J. C. Montero, T. R. Feldpausch, E. N. H. Coronado, T. J. Killeen, B. Mostacedo, R. Vasquez, R. L. Assis, J. Terborgh, F. Wittmann, A. Andrade, W. F. Laurance, S. G. W. Laurance, B. S. Marimon, B.-H. Marimon, I. C. Guimarães Vieira, I. L. Amaral, R. Brienen, H. Castellanos, D. Cárdenas López, J. F. Duivenvoorden, H. F. Mogollón, F. D. de A. Matos, N. Dávila, R. García-Villacorta, P. R. Stevenson Diaz, F. Costa, T. Emilio, C. Levis, J. Schietti, P. Souza, A. Alonso, F. Dallmeier, A. J. D. Montoya, M. T. Fernandez Piedade, A. Araujo-Murakami, L. Arroyo, R. Gribel, P. V. A. Fine, C. A. Peres, M. Toledo, G. A. Aymard C, T. R. Baker, C. Cerón, J. Engel, T. W. Henkel, P. Maas, P. Petronelli, J. Stropp, C. E. Zartman, D. Daly, D. Neill, M. Silveira, M. R. Paredes, J. Chave, D. de A. Lima Filho, P. M. Jørgensen, A. Fuentes, J. Schöngart, F. Cornejo Valverde, A. Di Fiore, E. M. Jimenez, M. C. Peñuela Mora, J. F. Phillips, G. Rivas, T. R. van Andel, P. von Hildebrand, B. Hoffman, E. L. Zent, Y. Malhi, A. Prieto, A. Rudas, A. R. Ruschell, N. Silva, V. Vos, S. Zent, A. A. Oliveira, A. C. Schutz, T. Gonzales, M. Trindade Nascimento, H. Ramirez-Angulo, R. Sierra, M. Tirado, M. N. Umaña Medina, G. van der Heijden, C. I. A. Vela, E. Vilanova Torre, C. Vriesendorp, O. Wang, K. R. Young, C. Baider, H. Balslev, C. Ferreira, I. Mesones, A. Torres-Lezama, L. E. Urrego Giraldo, R. Zagt, M. N. Alexiades, L. Hernandez, I. Huamantupa-Chuquimaco, W. Milliken, W. Palacios Cuenca, D. Pauletto, E. Valderrama Sandoval, L. Valenzuela Gamarra, K. G. Dexter, K. Feeley, G. Lopez-Gonzalez, M. R. Silman, S. P. Hubbell, F. He, R. Condit, L. Borda-de-Agua, J. Kellner, H. Ter Steege, G. A. Black, T. H. Dobzhansky, C. Pavan, J. M. Pires, T. Dobzhansky, G. A. Black, M. J. G. Hopkins, M. J. Costello, R. M. May, N. E. Stork, P. Haripersaud, H. ter Steege, J.-J. de Granville, H. Chevillotte, M. Hoff, D. P. Bebber, M. A. Carine, J. R. Wood, A. H. Wortley, D. J. Harris, G. T. Prance, G. Davidse, J. Paige, T. D. Pennington, N. K. Robson, R. W. Scotland, B. J. McGill, R. S. Etienne, J. S. Gray, D. Alonso, M. J. Anderson, H. K. Benecha, M. Dornelas, B. J. Enquist, J. L. Green, F. He, A. H. Hurlbert, A. E. Magurran, P. A. Marquet, B. A. Maurer, A. Ostling, C. U. Soykan, K. I. Ugland, E. P. White, R. J. Warren, D. K. Skelly, O. J. Schmitz, M. A. Bradford, N. C. A. Pitman, J. W. Terborgh, M. R. Silman, P. N. V, D. A. Neill, C. E. Cerón, W. A. Palacios, M. Aulestia, N. C. A. Pitman, M. R. Silman, J. W. Terborgh, F. D. Lozano, M. W. Schwartz, M. W. Schwartz, D. Simberloff, J. E. Richardson, R. T. Pennington, T. D. Pennington, P. M. Hollingsworth, T. L. Couvreur, F. Forest, W. J. Baker, S. Cavers, C. W. Dick, D. H. Janzen, S. A. Mangan, S. A. Schnitzer, E. A. Herre, K. M. Mack, M. C. Valencia, E. I. Sanchez, J. D. Bever, W. Balée, D. G. Campbell, C. Levis, P. F. de Souza, J. Schietti, T. Emilio, J. L. P. V. Pinto, C. R. Clement, F. R. C. Costa, D. A. Posey, E. Montoya, V. Rull, N. D. Stansell, M. B. Abbott, S. Nogué, B. W. Bird, W. A. Díaz, C. Gomez-Navarro, C. Jaramillo, F. Herrera, S. L. Wing, R. Callejas, C. H. McMichael, D. R. Piperno, M. B. Bush, M. R. Silman, A. R. Zimmerman, M. F. Raczka, L. C. Lobato, H. ter Steege, P. P. Haripersaud, O. S. Bánki, F. Schieving, S. J. Phillips, R. P. Anderson, R. E. Schapire, S. J. Phillips, M. Dudik, C. A. Quesada, J. Lloyd, L. O. Anderson, N. M. Fyllas, M. Schwarz, C. I. Czimczik, B. Rollet, H. ter Steege, N. C. Pitman, O. L. Phillips, J. Chave, D. Sabatier, A. Duque, J. F. Molino, M. F. Prévost, R. Spichiger, H. Castellanos, P. von Hildebrand, R. Vásquez, P. M. Fearnside, D. Mouillot, D. R. Bellwood, C. Baraloto, J. Chave, R. Galzin, M. Harmelin-Vivien, M. Kulbicki, S. Lavergne, S. Lavorel, N. Mouquet, C. E. Paine, J. Renaud, W. Thuiller, G. Lopez-Gonzalez, S. L. Lewis, M. Burkitt, O. L. Phillips, P. J. M. Maas, L. Y. T. Westra, H. Rainer, A. Q. Lobão, R. H. J. Erkens, P. V. A. Fine, D. C. Daly, G. V. Muñoz, I. Mesones, K. M. Cameron, K. J. Feeley, M. R. Silman, M. Dufrene, P. Legendre, J. Bunge, M. Fitzpatrick, J. Bunge, L. Woodard, D. Böhning, J. A. Foster, S. Connolly, H. K. Allen, J.-P. Z. Wang, B. G. Lindsay, A. Chao, R. K. Colwell, C.-W. Lin, N. J. Gotelli, C. X. Mao, R. K. Colwell, U. Brose, N. D. Martinez, R. J. Williams, I. J. Good, A. Chao, S.-M. Lee, I. Rocchetti, J. Bunge, D. Bohning, R. A. Fisher, A. S. Corbet, C. B. Williams, F. W. Preston, J.-P. Wang, P. V. A. Fine, and R. H. Ree, Hyperdominance in the Amazonian Tree Flora, Science 342, 1243092 (2013).

[13] L. H. M. Cosme, J. Schietti, F. R. C. Costa, and R. S. Oliveira, The Importance of Hydraulic Architecture to the Distribution Patterns of Trees in a Central Amazonian Forest, New Phytol. 215, 113 (2017).

[14] S. E. Russo, S. J. Davies, D. A. King, and S. Tan, Soil-Related Performance Variation and Distributions of Tree Species in a Bornean Rain Forest, J. Ecol. 93, 879 (2005).

[15] K. J. Gaston, Species-Range Size Distributions: Products of Speciation, Extinction and Transformation: Philosophical Transactions of the Royal Society of London Series B, 353, 219 (1998).

[16] R. E. Ricklefs, H. Qian, and P. S. White, The Region Effect on Mesoscale Plant Species Richness between Eastern Asia and Eastern North America, 2, 129 (2004).

[17] F. Munoz, B. R. Ramesh, and P. Couteron, How Do Habitat Filtering and Niche Conservatism Affect Community Composition at Different Taxonomic Resolutions?, Ecology 95, 2179 (2014).

[18] R. K. Colwell, C. Rahbek, and N. J. Gotelli, The Mid-Domain Effect and Species Richness Patterns:What Have We Learned so Far?, Am. Nat. 163, E1 (2004).

[19] H. ter Steege, N. Pitman, D. Sabatier, H. Castellanos, P. Van der Hout, D. C. Daly, M. Silveira, O. Phillips, R. Thomas, J. V. A. N. Essen, H. Mogollon, and W. Morawetz, A Spatial Model of Tree a -Diversity and Tree Density for the Amazon, Biodivers. Conserv. 12, 2255 (2003).

[20] T. F. Rangel, N. R. Edwards, P. B. Holden, J. A. F. Diniz-Filho, W. D. Gosling, M. T. P. Coelho, F. A. S. Cassemiro, C. Rahbek, and R. K. Colwell, Modeling the Ecology and Evolution of Biodiversity: Biogeographical Cradles, Museums, and Graves. Science, Science. 361, (2018).

